# A Flexible, Interpretable, and Accurate Approach for Imputing the Expression of Unmeasured Genes

**DOI:** 10.1101/2020.03.30.016675

**Authors:** Christopher A Mancuso, Jacob L Canfield, Deepak Singla, Arjun Krishnan

**Affiliations:** Department of Computational Mathematics, Science and Engineering, Michigan State University, East Lansing, MI 48824, USA; Department of Biochemistry and Molecular Biology, Michigan State University, East Lansing, MI 48824, USA; Indian Institute of Technology, Delhi, India.

## Abstract

While there are >2 million publicly-available human microarray gene-expression profiles, these profiles were measured using a variety of platforms that each cover a pre-defined, limited set of genes. Therefore, key to reanalyzing and integrating this massive data collection are methods that can computationally reconstitute the complete transcriptome in partially-measured microarray samples by imputing the expression of unmeasured genes. Current state-of-the-art imputation methods are tailored to samples from a specific platform and rely on gene-gene relationships regardless of the biological context of the target sample. We show that sparse regression models that capture sample-sample relationships (termed *SampleLASSO*), built on-the-fly for each new target sample to be imputed, outperform models based on fixed gene relationships. Extensive evaluation involving three machine learning algorithms (LASSO, k-nearest-neighbors, and deep-neural-networks), two gene subsets (GPL96-570 and LINCS), and three imputation tasks (within and across microarray/RNA-seq) establishes that *SampleLASSO* is the most accurate model. Additionally, we demonstrate the biological interpretability of this method by showing that, for imputing a target sample from a certain tissue, *SampleLASSO* automatically leverages training samples from the same tissue. Thus, *SampleLASSO* is a simple, yet powerful and flexible approach for harmonizing large-scale gene-expression data.

## Introduction

High-throughput gene expression technologies – especially microarray (Heller, 2002) and RNA-sequencing (RNA-seq) (Wang *et al.*, 2009) – have revolutionized our ability to capture and understand the large-scale cellular context of many biological systems in humans and several model organisms (Stark *et al.*, 2019; Hoheisel, 2006). Fortunately, due to community-wide norms and funding requirements, nearly all of the resulting transcriptomes have been deposited in publicly-available repositories (Lachmann *et al.*, 2018; Athar *et al.*, 2019; Edgar *et al.*, 2002; Barrett *et al.*, 2013). For example, as of 29 January 2020, there are >1 million human microarray samples from >24k datasets along with about half as much human RNA-seq data (>583k samples from ~12k datasets) contained in the NCBI Gene Expression Omnibus database (Edgar *et al.*, 2002; Barrett *et al.*, 2013).

The purpose of these publicly-available data is to enable other researchers to use published datasets to reproduce original findings, reuse datasets in new ways to answer new questions (Rung and Brazma, 2013), or combine thousands of datasets to build integrative models (Greene *et al.*, 2015) towards precision medicine (Alyass *et al.*, 2015). However, a major hurdle in realizing these goals is the fact that microarray profiles have been measured using a number of different platforms that each measure a different number of pre-defined genes (ranging from a few hundred genes to ~20k genes). For instance, the most popular genome-scale platform *Affymetrix Human Genome U133 Plus 2.0 Array* (GEO ID: *GPL570*) accounts for only 22% of the >1 million samples. The next most popular *Affymetrix Human Genome U133A Array* (GEO ID: *GPL96*) accounts for another 11% of the samples, but only covers <12k genes. Therefore, it is a significant challenge to gain insights about the full complement of genes in the human genome across the diversity of biological samples and unique experimental conditions in existing microarray data.

In addition to these researcher-submitted microarray datasets, concerted effort has also been put into defining a reduced set of genes that can be measured and then be used to accurately recover the expression of all the other genes (Donner *et al.*, 2012; Rudd *et al.*, 2015). The most prominent example of this effort is the Library of Integrated Network-Based Cellular Signatures (LINCS) microarray program (Subramanian *et al.*, 2017), which has shown that measuring 978 “landmark” genes, costing only $5 per sample (Peck *et al.*, 2006), is sufficient to then use to impute the expression of all other (tens of thousands of) genes. There are currently 1.3 million microarray samples in the LINCS data repository capturing the effect of numerous chemical and genetic perturbations on gene expression (Subramanian *et al.*, 2017).

With either of these massive data collections – the >1 million public transcriptomes from various microarray platforms or the 1.3 million LINCS profiles – restricting analysis and integration to the measured genes common to all platforms/samples will result in a tremendous loss of valuable data. Therefore, effectively leveraging the full data compendia on a genome-scale necessitates computational methods that can use the expression levels of the measured genes in a *partially-measured microarray sample* to impute the expression of all *unmeasured genes* in that sample to reconstitute a complete transcriptome [Fig. 1A]. A few previous studies have indeed proposed methods to solve this problem in various settings. Sparse regression models that use *gene-gene correlation signals* have been shown to be effective in imputing gene expression in samples from the GPL96 (<12k genes) microarray platform based on samples from the GPL570 (whole-genome) platform (Zhou *et al.*, 2017). Others have developed methods that use gene correlations based on low-rank regression (Ye *et al.*, 2013) and deep neural networks (Chen *et al.*, 2016; Wang *et al.*, 2018), specifically for the LINCS dataset. These methods rely on training machine learning models that map the relationship between fixed sets of measured and unmeasured genes in a specific setting, be it sparse gene-based regression for the GPL570-96 setting (Zhou *et al.*, 2017) or deep learning for the LINCS setting (Chen *et al.*, 2016; Wang *et al.*, 2018). Methods have also been proposed to address the problem of identifying the best reduced set of genes to measure to enable subsequent imputation of all other genes, again within the scope of specific large datasets (Donner *et al.*, 2012; Rudd *et al.*, 2015; Abid *et al.*, 2019). However, all these methods lack the flexibility for broad adoption since public datasets come from many different expression-profiling technologies, with each measuring the expression of a different subset of genes in the genome. All current methods are hard to adapt for imputing unmeasured gene-expression in an arbitrary experiment since they require training completely new models for every microarray platform (or every new reduced gene set design), which, in turn, requires very large datasets for model-training.

**Fig 1.**
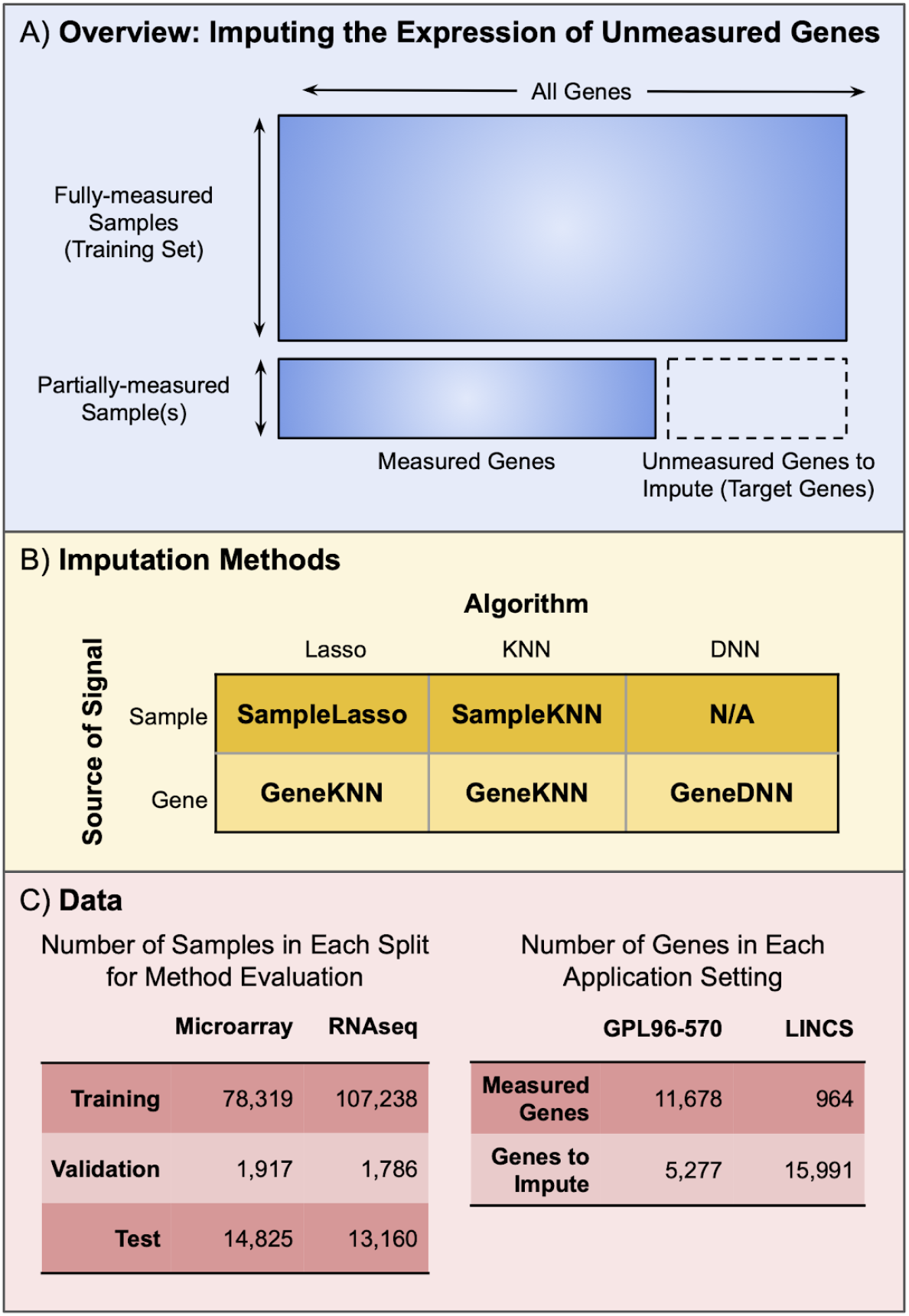
Overview of gene expression imputation. A) Schematic of the problem of “imputing the expression of unmeasured genes”. A training dataset is used to fill in the expression values of genes from partially-measured samples. The five methods (B) and summary of the data (C) used in this study. The methods are named using a combination of the machine learning algorithm and biological signal that the method uses.

Leveraging gene-gene correlations in data was an important component of gene expression imputation that focussed on the related-yet-distinct “missing value” problem [Fig. S1], concerned with recovering the expression values of individual genes that were lost *within a single dataset* due to arbitrary technical error in samples, i.e. filling arbitrary empty cells within a larger data matrix (Aittokallio, 2010; Brock *et al.*, 2008; Liew *et al.*, 2011). Many methods have been proposed to tackle this problem (Troyanskaya *et al.*, 2001; Bø *et al.*, 2004; Kim *et al.*, 2005; X. Wang *et al.*, 2006; Oba *et al.*, 2003; Kim *et al.*, 2004; Celton *et al.*, 2010), and, in general, imputing missing values has been shown to improve downstream tasks such as clustering, classification, co-expression network building, and differential expression (de Brevern *et al.*, 2004; Tuikkala *et al.*, 2008; D. Wang *et al.*, 2006; Oh *et al.*, 2011). Although these seminal works on the missing-value problem guide imputation methods today, the fact that we can now leverage information from >100k samples at a time to improve the imputation of unmeasured genes requires a rethinking of imputation strategies. Thus, it is critical that new imputation methods select only the most relevant samples to the target sample, as gene-gene correlations change across different biological contexts (Melé *et al.*, 2015).

In this study, we demonstrate that using a sparse-regression method that leverages information from the most similar samples provides more accurate predictions than other methods while also providing a highly interpretable underlying model. Current state-of-the-art imputation methods train machine learning models that capture the relationship of the predetermined set of measured genes to each predefined unmeasured gene (or set of unmeasured genes). Then, during imputation in an expression sample with the same measured/unmeasured genes, these methods use the pretrained models to impute the expression of the unmeasured genes. We propose a variant of this approach that we call *SampleLASSO* in which, for every new expression sample to be imputed, a new sparse regression model is trained on-the-fly that captures the relationship of this expression sample to all others in the training set based on the genes measured in that given sample. We compare our method to four other imputation methods based on three different algorithms – k-nearest neighbors, regularized linear regression, and deep neural networks – that leverage gene-gene or sample-sample relationships [Fig. 1B]. All these methods are evaluated on imputation within the same-technology (microarray and RNA-seq), as well as using RNA-seq data to impute microarray data. The evaluation is carried out for two different, practical unmeasured gene settings: 1) a high number of measured genes and low number of unmeasured genes (GPL96-570 gene subset) and 2) a small number of measured genes and large number of unmeasured genes (LINCS gene subset) [Fig. 1C]. Extensive evaluations using multiple accuracy metrics within a rigorous temporally-split and dataset-preserving scheme showed that, for all cases considered, the flexible sample-based imputation (*SampleLASSO)* is always the best performing method. We also demonstrate the biological interpretability of this method by showing that, for imputing a given sample from a certain tissue, the *SampleLASSO* model automatically up-weights training samples from the same tissue type.

## Materials and Methods

### Data

We used gene expression data from both microarray and RNA-seq technologies. For the microarray data, we downloaded all human samples from the *Affymetrix Human Genome U133 Plus 2.0 Array* (also known as the GPL570 platform) for NCBI GEO (Barrett *et al.*, 2013) as raw CEL files and performed background subtraction, quantile transformation, and summarization using fRMA (McCall *et al.*, 2010) based on a custom CDF (Dai *et al.*, 2005) mapping probes to Entrez gene IDs. This yielded 108,205 samples with 19,702 genes. For the RNA-seq data, we downloaded all 133,776 human, TPM-normalized samples from ARCHS4 (Lachmann *et al.*, 2018), and further processed the data by converting ENST IDs to Entrez gene IDs for only the genes found in the microarray data. Genes that could not be mapped this way were discarded from both microarray and RNA-seq data. This yielded a total of 16,955 genes. Finally, the RNA-seq data was then arcsinh transformed. More information on data processing is provided in Section 1.1 of the Supplemental Material.

### Validation Scheme

#### Subsetting Genes

To evaluate the imputation methods, we chose to split genes into measured and unmeasured sets to represent two very different practical scenarios [Fig. 1C]. First, we considered the situation in which we have a large number of measured genes that we could use to impute a smaller number of unmeasured genes. This scenario presents itself in the problem of using the 11,678 genes measured in the older human microarray platform *Affymetrix Human Genome U133A Array* (also known as GPL96) to then impute the expression of an additional 5,277 genes that are only present in the newer genome-scale platform *Affymetrix Human Genome U133 Plus 2.0 Array* (also known as GPL570) (Zhou *et al.*, 2017). This gene-split is referred to as the *GPL96-570 gene subset* in this work. Second, we considered the situation in which we have a small number of measured genes that we could use to impute a large number of unmeasured genes. For this scenario we used 964 “landmark” genes from LINCS as the measured genes to impute the expression of all the other genes in the genome-scale *Affymetrix Human Genome U133 Plus 2.0 Array* (15,991 unmeasured genes) (Subramanian *et al.*, 2017; Chen *et al.*, 2016). This gene-split is referred to as the *LINCS gene subset* in this work.

#### Splitting Samples

We divided the expression samples into training, validation, and testing sets. The training data was used to fit the models, the validation data was used for hyperparameter tuning, and the testing data was used in the final evaluations of the models (shown in all the figures in the main text). To mitigate data leakage, we ensured that entire datasets were assigned to splits, thus keeping all expression samples from the same experiment (dataset) together in the same split. The data was also temporally split, with the oldest expression samples being placed in the training and validation sets, and the newest samples going into the test set. To speed up hyperparameter tuning, which consisted of training >500,000 individual models, we further subsetted the validation set by taking 10% of the expression samples from each experiment in the full validation set (or at least 2 expression samples, if the number of samples in an experiment was less than 20) (see Section 1.1 in Supplemental Material and Fig. S2).

For all imputation methods, we standardized each feature in the training set by subtracting the mean and dividing by the standard deviation of the given feature. Correspondingly, each feature in the validation and test sets was standardized using the mean and standard deviation obtained from the training set.

### Imputation Methods

In this study, we evaluated imputation methods using three distinct machine-learning algorithms: *k*-nearest neighbors (*KNN*), least absolute shrinkage and selection operator (*LASSO)*, and deep neural networks (*DNN*).

*KNN* is a machine learning algorithm that predicts the target variable for every new example based on the target variables of the *k* most similar examples in the training data. To impute a given target variable, we used a weighted average of the measured target variable from the *k* most similar examples based on Euclidean distance, with the weight equal to the inverse of the distance. KNN imputation is a widely-used imputation method for gene expression and provides a strong baseline (Troyanskaya *et al.*, 2001; Brock *et al.*, 2008; Donner *et al.*, 2012; Chen *et al.*, 2016).

*LASSO* is a linear regression method in the family of least-squares optimizers (Tibshirani, 1996). LASSO builds a sparse model for a given target variable using the following cost function:

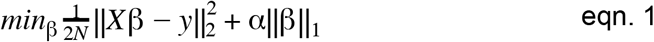

where β is the vector of learned parameters, *X* is the training data, *y* is the target variable, α is the hyperparameter that determines the extent of L1-regularization (||β||_1_). L1-regularization prevents overfitting by setting many of the elements of β to 0. While a variety of least-squares optimizers have been applied to the gene expression imputation problem (Bø *et al.*, 2004; Nguyen *et al.*, 2004; Hu *et al.*, 2006; Brock *et al.*, 2008), LASSO is the method most suited when the number of features is large (Zhang and Huang, 2008; Chen *et al.*, 2016; Donner *et al.*, 2012). In this study, we used the KNN and LASSO implementations contained in the Python package *scikit-learn* (Pedregosa *et al.*, 2011).

*DNN* is a multi-layer neural network with bespoke architectures designed for each machine learning task. We used a multi-layered feed-forward neural network, similar to the model used in a previous work that evaluated the feasibility of using DNNs for imputing gene expressions using the LINCS landmark genes (Chen *et al.*, 2016). DNN models were trained using Nividia Tesla k80 GPUs and implemented using the Python package *Keras* (Chollet, 2015) with a *Tensorflow* backend (Abadi *et al.*, 2016). To most accurately mimic the DNN architecture presented in (Chen *et al.*, 2016), we randomly split the unmeasured genes into four sets (3998 genes per set) and trained a DNN for each set. For more information on the DNN method see Section 1.2 of the Supplemental Material.

We used these three algorithms – KNN, LASSO, DNN – to leverage two distinct types signals – gene-gene similarities (across samples) and sample-sample similarities (across genes) – for imputing the expression of unmeasured genes in a new partially-measured sample, resulting in five methods referred to in this study as *SampleKNN*, *GeneKNN*, *SampleLASSO* (proposed here), *GeneLASSO*, and *GeneDNN*. For intuitive, pictorial schematics of the five methods, see Figs. S3-S7 in Section 1.3 of the Supplemental Material.

*SampleKNN* and *GeneKNN* are the most straightforward and popular implementation of KNN for gene expression imputation. For a new partially-measured expression sample to be imputed, *SampleKNN* works by first finding the (*k*) most similar samples in the training set based on the expression of all measured genes, and then imputing the expression of each unmeasured gene with the weighted average of that gene’s expression in the most similar training samples. Thus, the major biological signal used is the similarity between samples (across genes). Conversely, *GeneKNN* works on a gene-by-gene basis. For each gene that is missing (unmeasured) in a new sample, the method first finds the (*k*) measured genes most similar in their expression pattern across all the samples in the training set, and then imputes the expression of the unmeasured gene with the weighted average of the expression of those *k* genes in that new sample. Thus, the major biological signal used is the similarity between genes (across samples).

*GeneLASSO* is the traditional, widely-adopted means of implementing LASSO for gene expression imputation. Here, using the fully-measured training set, a separate sparse regression model is trained for each unmeasured gene, to predict its expression based on a linear combination of all the measured genes. Then, given a new partially-measured sample, the expression of every unmeasured gene is imputed using that gene’s pre-trained model, with the predicted expression being equal to the sum of the expression of the measured genes in the new sample weighted by the model coefficients. Akin to *GeneKNN*, the main source of biological signal for *GeneLASSO* is gene-gene expression similarities. As an alternative to *GeneLASSO*, which requires information about which genes are unmeasured in a new sample and a pre-trained model for each of those genes, in this study, we propose a simple alternative called *SampleLASSO*. Given a new partially-measured sample, *SampleLASSO* builds a single model on-the-fly that predicts that sample’s expression profile based on a sparse linear combination of all the samples in the training set only using the subset of genes measured in the new sample. Here, for every sample to be imputed, the coefficients of the trained model in essence finds the relationship of that sample to all samples in the training set. Then, all the unmeasured genes are imputed using this trained sample-specific model. The main source of biological signal in SampleLASSO, thus, comes from sample similarities. We note a method similar to *SampleLASSO* has been reported before (called *LS_array*) (Bø *et al.*, 2004). However, the implementation of that method was focused on the missing value problem, and has never been applied to the big-data and the unmeasured gene problem.

*GeneDNN* uses a DNN to predict the expression of (a fixed set of) many unmeasured genes using a single model that captures a complex, nonlinear relationship between the unmeasured genes and the measured genes using the training set. Thus, the main biological signal in *GeneDNN* is also from gene-gene relationships. Since DNNs are significantly harder to build, train, and parameterize than LASSO and KNN methods, we only considered DNN on the LINCS gene subset as the optimal model architecture was worked out in (Chen *et al.*, 2016).

### Hyperparameter Tuning

For both KNN and LASSO, there is only one hyperparameter that needs to be tuned; *k:* the number of most similar training examples to consider in KNN and α: the parameter that sets the strength of the L1-regularization term in LASSO. Although there are many hyperparameters to tune in a DNN, we fixed most parameters based on optimal values found in (Chen *et al.*, 2016), and just tuned the optimizer and learning rate. Hyperparameter tuning was done using the validation set data [Section 1.4 in Supplemental Material, Fig. S8-S12, and Table S1], and the optimal hyperparameters were then used for final evaluation using the test set. We note that for *GeneDNN* we used all the ~13k samples in the validation set for hyperparameter tuning as: 1) this most closely mimics the situation in (Chen *et al.*, 2016), and 2) DNNs additionally use the validation to determine which epoch (i.e. how many passes through the data the model goes through) yields the best model.

### Evaluation Metrics

A commonly used metric for evaluating gene expression imputation methods is Normalized Root Mean Square Error (NRMSE). The NRMSE for a gene (*g*_*i*_) is given by:

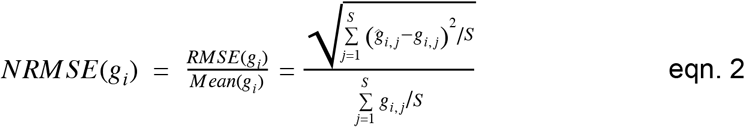

where *RMSE* is the root mean square error, *S* is the number of samples, and 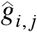, *g*_*i, j*_ are the imputed and real expression values, respectively, for the *i*^*th*^ gene in the *j*^*th*^ sample.

In addition to NRMSE, we also report evaluation results using the Spearman correlation coefficient and the Mean Absolute Error (MAE) in Section 2.2 of the Supplemental Material.

### Interpreting *SampleLASSO* Models

We evaluated the interpretability of *SampleLASSO* models by examining if the β -coefficients of a model trained for a particular sample recapitulated that sample’s tissue-of-origin by assigning high positive *β* values to samples in the training set from the same tissue relative to samples from all other tissues.

Specifically, for a given target sample *s* that we built a *SampleLASSO* model for, we calculated a z-score, *z*_*s, T*_, for each tissue *T* in this annotated set based on the β values of training samples from that tissue:

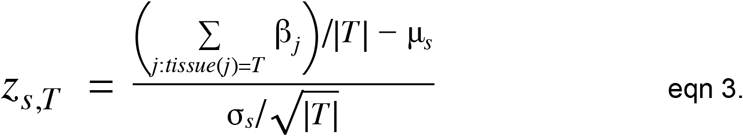

where |*T*| is the number of labeled samples for tissue *T*, β_*j*_ is the value of the β -coefficient from the *SampleLASSO* model for the *j*-th sample, and μ_*s*_ and σ_*s*_ are the mean and standard deviation of the β -coefficients of all samples in the training set that have any tissue label.

To perform this analysis we used a large set of expression samples that were manually-curated to their tissue-of-origin (Lee *et al.*, 2013). However, due to the initial temporal split of the data into training, validation, and test sets, all the labeled expression samples were in the original training set. Hence, just for this interpretability analysis, we separated out a subset of the tissue-labelled samples in the original training set into a new manually-curated test set. We created this subset so that it spanned six tissues that were sufficiently diverse and were labeled to at least 10 samples from at least 3 different datasets in both the training and the test sets. This resulted in the new test set having 222 expression samples from 29 different datasets and the new training set having 4,397 expression samples from 120 different datasets [Table. S2]. To calculate μ_*s*_ and σ_*s*_ we used any sample that had a tissue label, regardless of tissue type, allowing us to use 11,618 samples for these calculations. A *SampleLASSO* model was trained for each manually labeled test sample and used in the z-score analysis above (eqn. 3).

## Results

In this study, we compare imputation methods that use three distinct machine learning algorithms: least absolute shrinkage and selection operator (LASSO), k-Nearest Neighbors (KNN) and deep neural network (DNN) [Fig. 1B]. Combining these algorithms with the source of the data signal – gene-gene or sample-sample relationships – resulted in five imputation methods: *SampleLASSO*, *GeneLASSO*, *SampleKNN*, *GeneKNN*, and *GeneDNN*. These methods are evaluated in two settings with different sets of unmeasured genes [Fig. 1C]: 1) the GPL96-570 gene subset, which uses a relatively large number of genes (~11,000) to impute the expression of a smaller number of genes (~5,000) and 2) the LINCS gene subset, which uses a relatively small number of genes (~1,000) to impute the expression of a large number of genes (~16,000). We consider imputation using data from the same technology (using microarray to impute microarray, and RNA-seq to impute RNA-seq) as well as across technologies (using RNA-seq to impute microarray data). We evaluate methods using both the scale-free regression error metric (normalized root mean squared error; NRMSE), as well as Spearman correlation and mean absolute error. Lastly, we examine the model coefficients learned by *SampleLASSO* for biological interpretability.

We first evaluated the performance of the five imputation techniques using microarray data to impute microarray data [Fig. 2]. For both gene subset tasks, *SampleLASSO* is the best performing model, with both KNN methods performing relatively poorly. For the GPL96-570 gene split, *SampleLASSO* outperforms *GeneLASSO* 91% of the time, *SampleKNN* 99% of the time, and *GeneKNN* 100% of the time. For the LINCS gene split, these percentages are 90%, 98%, and 100%, respectively. *SampleLASSO* outperforms *GeneDNN*, the current best method for imputing unmeasured genes using LINCS landmark genes (Chen *et al.*, 2016), 76% of the time. Statistical tests further confirmed that SampleLASSO is significantly more accurate than all the other methods (p-value << 1E-3; paired Wilcoxon ranked-sum test). Table S2 provides a tabular form of this information. We find similar results when measuring imputation accuracy using Spearman correlation and mean absolute error [Figs. S13-S15].

**Fig 2.**
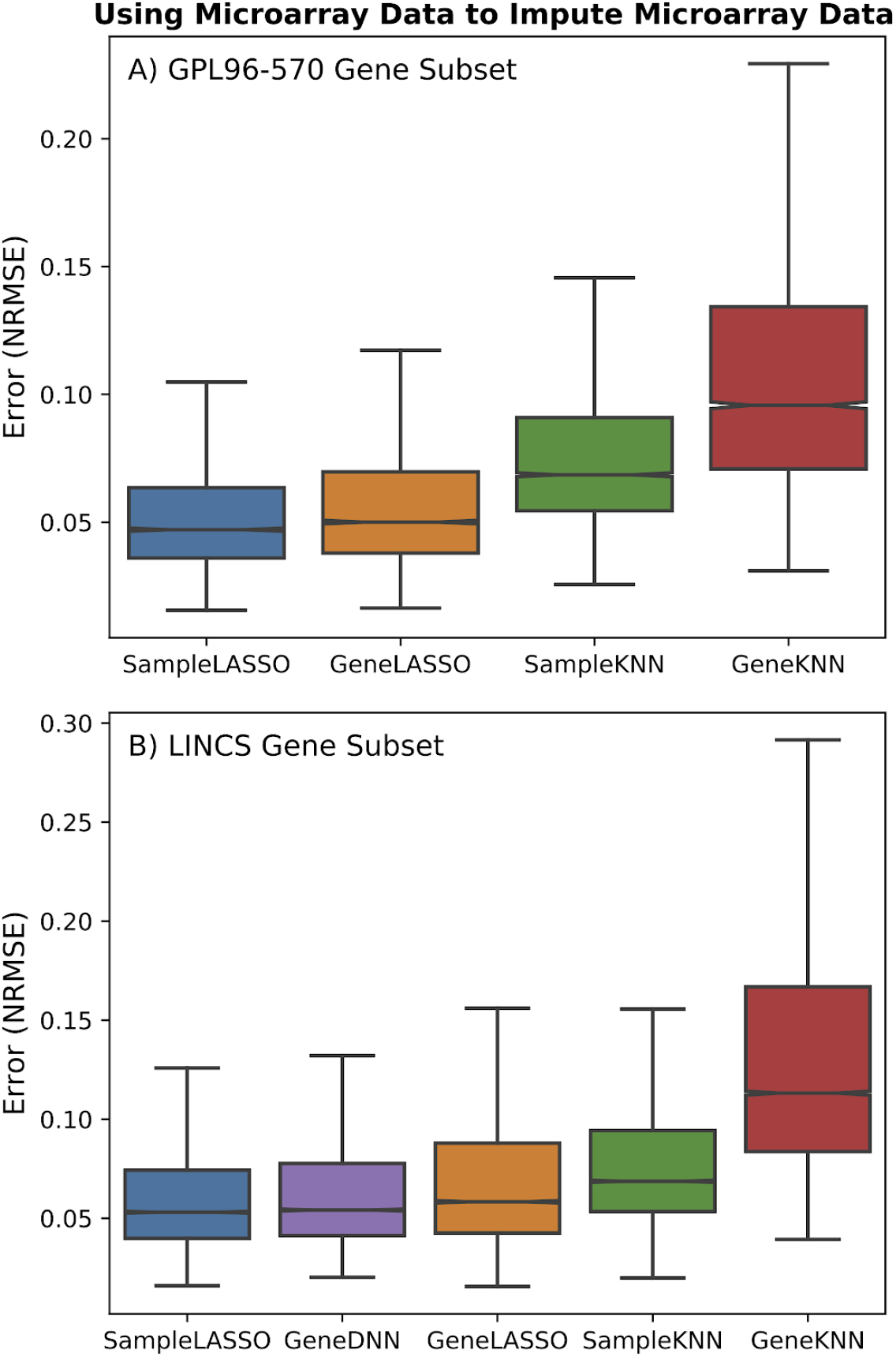
Performance of imputation methods for microarray data. Boxplots showing the performance of the five imputation methods (*SampleLASSO*, *GeneDNN*, *GeneLASSO*, *SampleKNN*, *GeneKNN*) across two gene subsets (A: GPL96-570 and B: LINCS), trained and imputed on microarray data. The evaluation metric is NRMSE, with lower values indicating better performance. The results show that *SampleLASSO* is the best performing method in both cases.

Although microarray platforms like the *Affymetrix Human Genome U133 Plus 2.0 Array* are able to quantify the expression of nearly all protein-coding genes, RNA-seq technology enables the quantification of nearly all cellular transcripts from both annotated and unannotated genes. Hence, it would be valuable to use RNA-seq data to predict the expression of genes missing in microarrays, enabling 1) re-analysis of novel genes in experimental settings captured in the vast number of microarray datasets and 2) joint analysis and integration of RNA-seq and microarray data based on a common set of genes. We evaluated the performance of using RNA-seq data to impute microarray data using the GPL96-570 and LINCS gene subsets [Fig. 3]. *SampleLASSO* is again the best performing method for both gene subsets. While *GeneLASSO* is the second best performing model in the LINCS gene subset, it is outperformed by *GeneKNN* in the GPL96-570 gene subset. *SampleKNN* is the worst performing method in both gene subsets. For the GPL96-570 gene split, *SampleLASSO* outperforms *GeneLASSO* 71% of the time, *SampleKNN* 80% of the time, and *GeneKNN* 70% of the time. For the LINCS gene split, these percentages change to 62%, 76% and 84%, respectively. *SampleLASSO* also outperforms *GeneDNN* 77% of the time. The Wilcoxon ranked-sum test between *SampleLASSO* and the other methods showed the performance increase of *SampleLASSO* was always statistically significant (p-value << 0.001). For the sake of completion, we also tested using RNA-seq data to impute RNA-seq data – and found that, with the exception of *GeneDNN* for the LINCS gene subset, *SampleLASSO* is again the best method [Fig. S16]. We note that using RNA-seq data to impute RNA-seq data is not a practically important task, as RNA-seq technologies do not require pre-determining a set of genes to measure.

**Fig 3.**
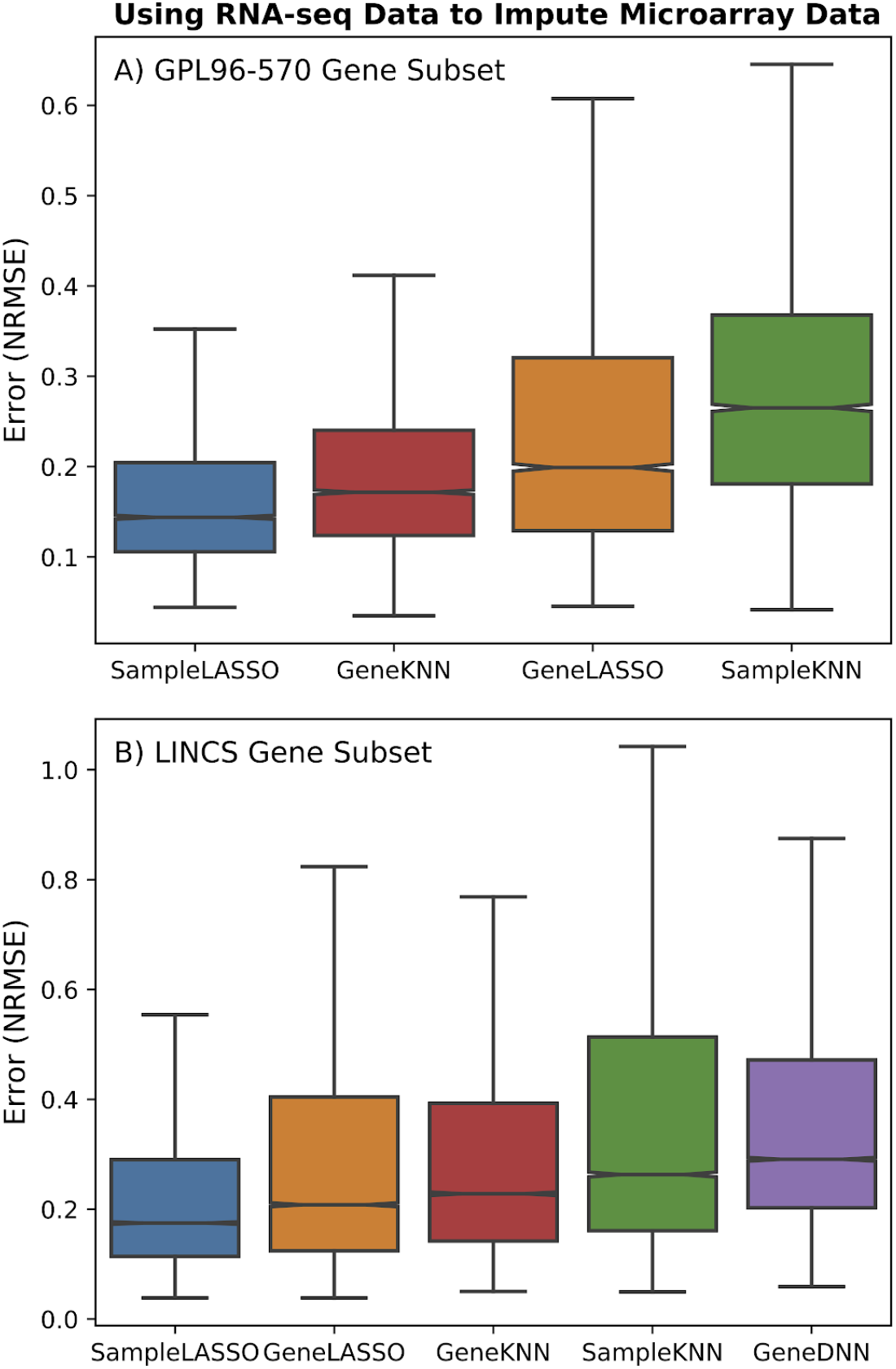
Performance of imputation methods for cross-technology imputation. Boxplots showing the performance of the five imputation methods (*SampleLASSO*, *GeneDNN*, *GeneLASSO*, *SampleKNN*, *GeneKNN*) across two gene subsets (A: GPL96-570 and B: LINCS) using RNA-seq data to impute microarray data. The evaluation metric is NRMSE, with lower values indicating better performance. The results show that *SampleLASSO* is the best performing method in both cases.

We additionally looked at the performance of the imputation methods as a function of the mean expression and variance of the unmeasured genes. For each unmeasured gene, we placed it into a low, medium or high bin based on its mean expression and variance in the context of the known expression values across all the samples [Fig. S17-S22]. Although the relative performance of the imputation methods doesn’t change too much when looking at the different categories of mean expression and variance, it can be seen in some cases that *SampleLASSO* performs particularly well for genes with low mean expression and high variance, which is the hardest category of genes to impute.

Although not very accurate in imputing unmeasured genes, a method such as *SampleKNN* offers immediate interpretability via the biological/experimental contexts of the *k*-nearest training samples picked by the method. Since we devised *SampleLASSO* with this desirable property in mind, we tested if this new method also offers biological interpretability in addition to providing very accurate imputation. Since gene expression samples have clear signals pertaining to their tissue-of-origin (Melé *et al.*, 2015), we focused on testing if the *SampleLASSO* model trained for a new sample from a particular tissue up-weighted training samples from that same tissue relative to training samples from other tissues.

To perform this analysis we used a large set of expression samples that were manually labeled to their tissue-of-origin (Lee *et al.*, 2013). Due to limited labelled tissues and their representation in our data, the analysis was restricted to six sufficiently-diverse tissues that had at least 10 samples from at least 3 different datasets in both the training set (total of 4,397 samples, 120 datasets) and the test set (total of 222 samples, 29 datasets; Table S3). Then, for each test sample, we trained a SampleLASSO model and used the model coefficients (β -coefficients) to calculate a z-score for each of the six tissues that represents the aggregate β -coefficients corresponding to training samples just from that tissue relative to the background distribution of β -coefficients of all labeled training samples (not just samples labeled from the six tissues). Thus, a large positive z-score for a particular tissue means that samples from that tissue were more informative than others. See *Materials and Methods* for more details. Implementing this analysis on the entire labeled test set, we observe that for most test samples, the strongest signal captured by *SampleLASSO* comes from training samples from the same tissue as the sample being imputed [Fig. 4, Fig. S23].

**Fig 4.**
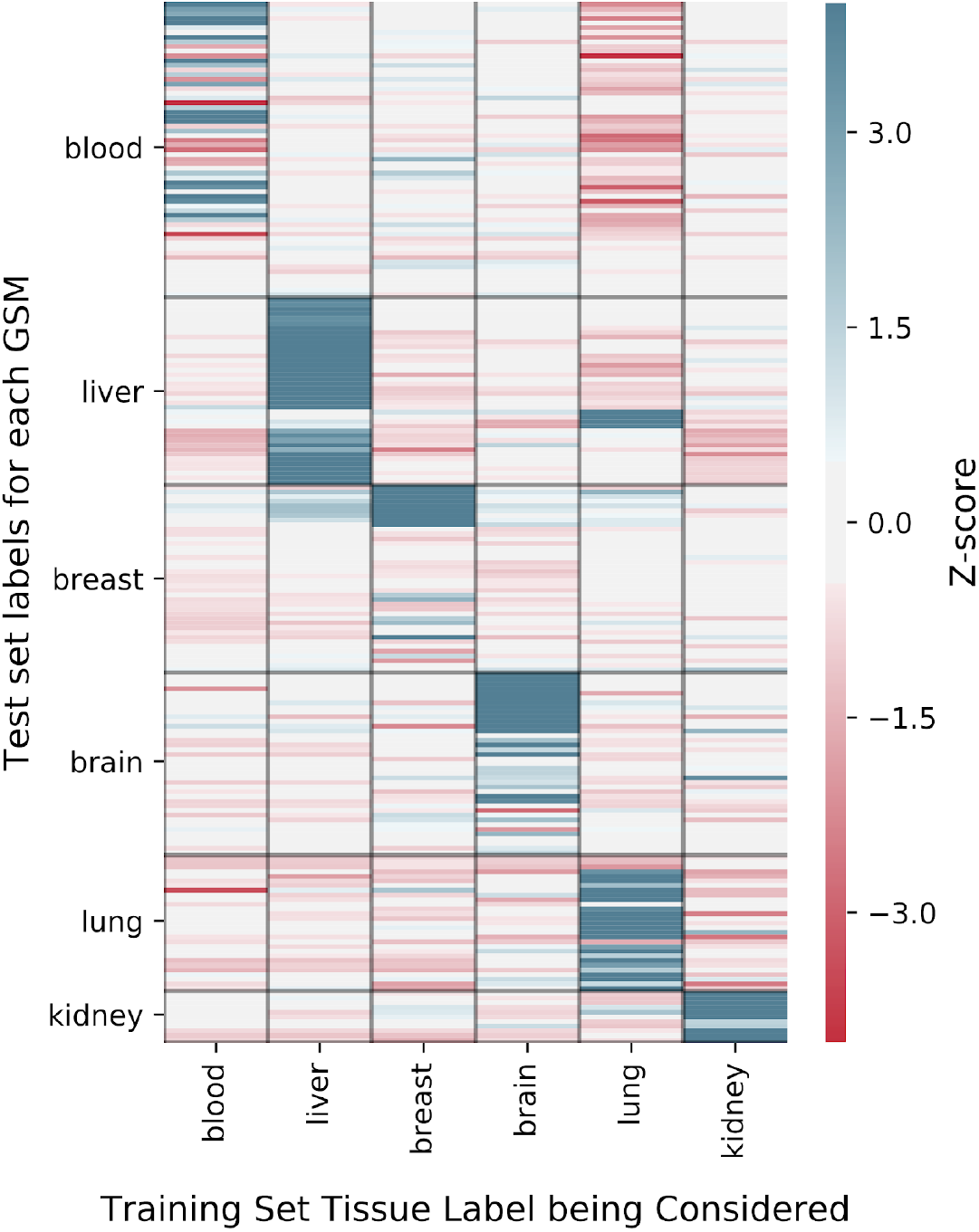
*SampleLASSO* captures biologically relevant information. Biological interpretability was evaluated using 4,619 expression samples (from 149 different experiments) labeled with tissue-of-origin to determine if *SampleLASSO* up-weights training samples of the sam tissue as the test sample in the sparse model. The rows represent the 222 tissue-labeled samples (GSMs) in the test set. The black horizontal lines separate test samples from different tissues. The columns correspond to the tissue type of the training samples. The colors of the heatmap represent the z-scores. Calculated per test sample (row), the z-score per tissue (column) corresponds to the normalized aggregate of the model coefficients of all the training samples from that tissue. See *Material and Methods* for more details. The diagonal blocks correspond to the case where the z-scores were calculated for the same tissue type as the tissue-of-origin of the test sample.

To aid in reproducibility, we have publicly released all the processed data that we used on Zenodo (https://doi.org/10.5281/zenodo.3711089) as well as all the code to re-generate the results and figures on GitHub (https://github.com/krishnanlab/Expresto). In addition, we also provide a user-friendly function that performs imputation using *SampleLASSO* given a file of expression data. The function will return the imputed expression values as well as a list of the most utilized samples from the training set (see Table S4 for an example).

## Discussion

In this study, we propose a simple, new method termed *SampleLASSO* for imputing the expression of unmeasured genes in partially-measured gene-expression profiles. *SampleLASSO* is a sparse regression model that trains a machine learning model on-the-fly for every expression profile that needs to have expression values imputed. Our extensive evaluations demonstrate that *SampleLASSO* outperforms all the other methods – consistently in a statistically significant manner – based on multiple error metrics, uniformly for unmeasured genes with a broad range of means and variances, in three different imputation tasks (within and across technologies), and in two imputation settings that differ in the number of measured genes by an order of magnitude. *SampleLASSO*’s strength comes from its ability to effectively leverage information from samples from the same biological context.

In addition to helping estimate the performance of imputation methods, our analyses in different imputation settings highlight various data standardization/normalization scenarios. When using microarray data to impute microarray data, the training data and validation/testing data (including both measured and unmeasured genes) are quantile normalized to the same distribution. When using RNA-seq data to impute RNA-seq data, samples only undergo within-sample normalization (using TPM) without any between-sample normalization. When using RNA-seq data to impute microarray, the training set data is not jointly normalized but the validation/testing data are. The fact that microarray data has a much lower imputation error than the RNA-seq also points to RNA-seq profiles having very different data distributions due to not being (quantile) normalized across samples, coming from many different sequencing platforms, and having a broader dynamic range than microarray data. Future work is required to examine the effect of data normalization and transformation and to develop strategies to perhaps transform data just based on measured genes and, upon imputation, recast the unmeasured genes into the original data space.

The performance of the various imputation methods in the cross-technology imputation task, where the influence of data transformation is most evident, highlights how each method works to impute gene expression. *SampleKNN* performs poorly because, for a given microarray sample to impute, it finds the closest RNA-seq samples, which come from a different data distribution. On the other hand, *GeneKNN* has relatively good performance because it works completely within the training data (RNA-seq) to find the genes nearest to a particular gene and then uses these gene relationships for imputation within the microarray data. Even though *GeneLASSO* similarly captures gene relationships only using the training data (RNA-seq), the mapping in the form of model coefficients does not transfer to microarray samples as easily as nearest neighbors. Similar issues, in addition to potential overfitting to the training set, thwart the performance of *GeneDNN* [Figs. S24-S26]. *SampleLASSO*’s top performance stems from having the unique property of learning a supervised model that, in addition to learning meaningful sample relationships, naturally captures the scaling factors required to closely map RNA-seq data distribution to microarray data distribution.

Although DNNs have been shown to give promising results in gene expression imputation in the LINCS setting (Chen *et al.*, 2016; Wang *et al.*, 2018), using them for imputation has a number of practical drawbacks. The foremost limitation is that a very large number of hyperparameters, in addition to other aspects of model optimization such as regularization and dropout, need to be tuned to build an accurate model. Next, complex models like DNNs are likely to overfit to the training set, as can be seen in the cross-technology setting (very different training and test data) where the drop in performance of *GeneDNN* is more severe than that of *GeneLASSO* on an independent set [Fig. 3, Figs. S24-S26]. DNNs are also hard to scale in terms of model size, with the number of weights growing nonlinearly with increasing layer number and layer size. Additionally, the current state of the standard hardware is such that only ~32GB of a model can fit into the memory of a GPU, and anything more requires the utilization of multiple GPUs.

In contrast, *SampleLASSO* is a simple, intuitive, flexible model. A number of the benefits of *SampleLASSO* emanate from the fact that a new machine learning model is trained on-the-fly for each new target sample that needs to be imputed based on the set of measured genes in that sample. Any set of genes can be measured/unmeasured in this setup, obviating the need for fixed pretrained models. Target data can also remain in the original scale/space without the need for data transformation. These benefits come along with *SampleLASSO*’s ability to leverage biological information specific to the new target sample, enabling easy interpretability. We specifically evaluated *SampleLASSO*’s interpretability on a large-scale using 4,397 samples labelled to various tissues of origin. This analysis shows that, when imputing a sample from a specific tissue, the *SampleLASSO* model up-weights training samples from the same tissue in majority of the cases [Fig. 4 and Fig. S23]. The rest of the cases could be due to high tissue heterogeneity (as in blood) or factors other than tissue-type (e.g. disease status, drug dosage) being the dominant signal.

In conclusion, we propose *SampleLASSO*, a simple method for imputing the expression of unmeasured genes. Extensive evaluations and analyses demonstrate that *SampleLASSO* is accurate, flexible, and interpretable. We have made all the data and code from this study freely available on Zenodo and GitHub (https://github.com/krishnanlab/Expresto) to aid in reproducing all our findings. Using a convenient function in our code, researchers can also use *SampleLASSO* to readily impute unmeasured genes in their samples of interest in any of the following practical settings: 1) Complete the expression profile of publicly-available microarray samples from any platform to make them comparable to the human whole-genome microarray, 2) Predict the expression of genes absent in standard microarrays (e.g. most non-protein coding genes) using RNA-seq to impute microarray samples, and 3) Fill in and effectively use genome-scale chemical and genetic perturbation expression data from LINCS based on the measured landmark genes.

## Acknowledgements

We thank Kayla A. Johnson, Nathaniel Hawkins, and the rest of the Krishnan Lab for valuable discussions and feedback on the manuscript.

