## Supplemental Material for "A Flexible, Interpretable, and Accurate Approach for Imputing the Expression of Unmeasured Genes"

#### Section 1: Supplemental Material for Methods

##### Section 1.1: Pictorial Representation of Unmeasured Gene Imputation

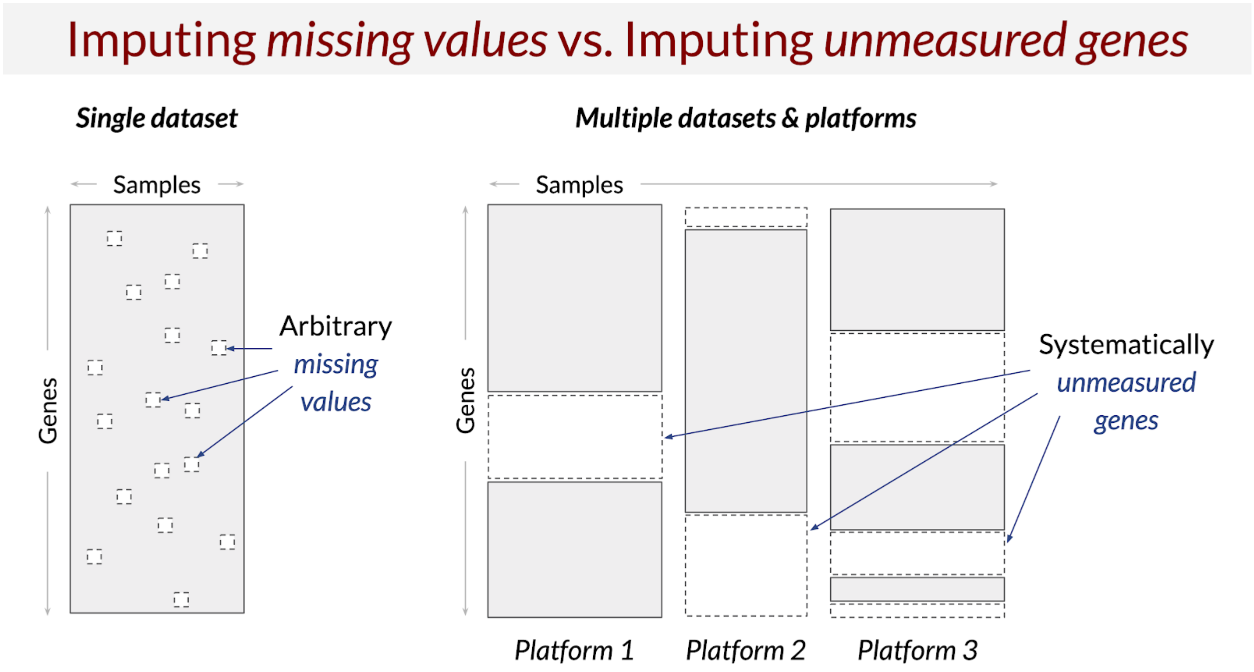


**Fig. S1. Schematic of the difference between "missing value" imputation versus "unmeasured gene" imputation.** In the missing value imputation problem, gene expression values from a single dataset that were unmeasured due to technical errors are imputed. In the unmeasured gene imputation problem, genes not measured at all by a given platform are imputed.

##### Section 1.2: Description of data processing steps

We downloaded the microarray data on Dec 6th 2017, and downloaded the RNA-seq data from the Sept 14th 2018 release of ARCHS4. In the microarray data, log transformation and quantile normalization was performed using Frozen Robust Multi-array Analysis (McCall *et al.*, 2010) and the probes were mapped to Entrez space using a custom CDF (Dai *et al.*, 2005). For the RNA-seq data, we converted the ENST IDs to Entrez IDs by taking the sum of all ENSTs mapped to a given ENSG, where the ENST to ENSG mapping was given by using the *gene2ensembl.gz* file available on the NCBI website on June 11th 2019. We note that the microarray data can be readily quantile normalized as all the data comes from the *Affymetrix Human Genome U133 Plus 2.0 Array* platform. In contrast, quantile normalization of the RNA-seq data is not straightforward as the data was generated using many different sequencing platforms, and it is difficult to perform batch effect correction across this large number of expression samples. We mention that although there are 984 “landmark” genes in original LINCS data, the software package we used to map from microarray probes to Entrez gene IDs only contained 964 of the LINCS genes.

The procedure used to split the expression samples into the training, validation and test sets was; 1) samples were first grouped together if they appear in the same experiments, 2) a date is assigned to every one of these experiment groups by selecting the oldest date of an expression sample associated within the group, and 3) the experimental groups are then temporally split into training, validation and testing sets with the oldest experimental groups going into the training set and most recent experimental groups going into the test set. The split ratios were such that we had 80% of the data in the training, 10% of the data in the validation and testing sets. We then further subset the validation to 10% of it’s full size as described in the main text. We note, for the DNN method we used all the validation data to most closely replicate the methods reported in (Chen *et al.*, 2016).

To get an idea if subsetting the validation data was sufficient we plotted the results for both the validation and test test for each method using the optimal hyperparameter [Fig. S2]. For both the Microarray-Microarray and RNAseq-Microarray cases, the performance between the two sets is nearly identical. For the situation where the validation and test data is RNA-seq (RNAseq-RNAseq), the test set performance is slightly worse than the validation set. However, we do not believe this is due to too few validation samples, rather it is due to the fact that since the data is split temporally, this is a batch effect as the sequencing technology is changing over time.

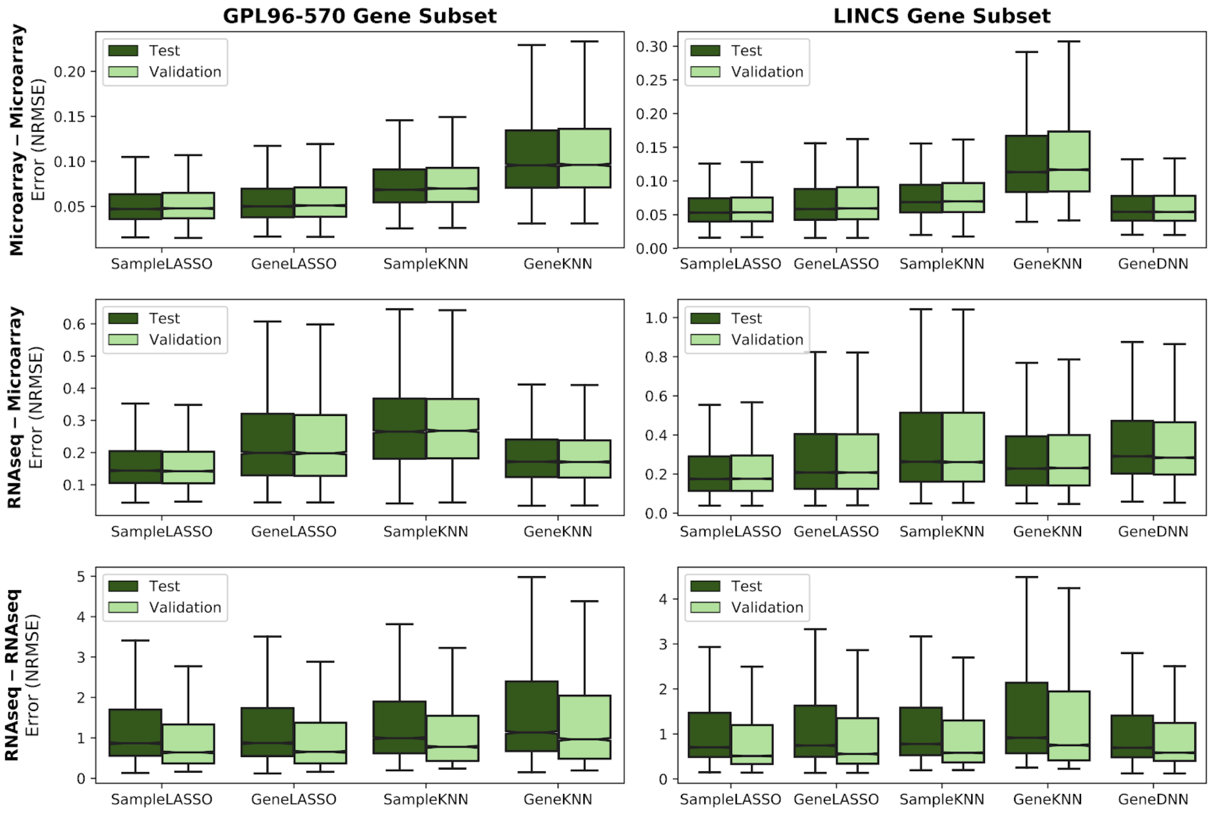


**Fig. S2. Comparison of imputation methods across both validation and test sets.** The performance of the methods is compared for the validation and test sets to see if using the subset validation set was sufficient. When the validation and testing data is microarray (top and middle rows) the performance is nearly identical. When the validation and test data is RNA-seq (bottom row) the test set performance is slightly worse than the validation set, but this effect is mostly likely due to the validation and test set containing expression samples from different sequencing platforms.

#### Section 1.3: Description of imputation methods

In this section, Figs. S3-S7 provide a pictorial representation of how each imputation method was implemented. Following these figures is a description of all the parameters that were used for the GeneDNN method.

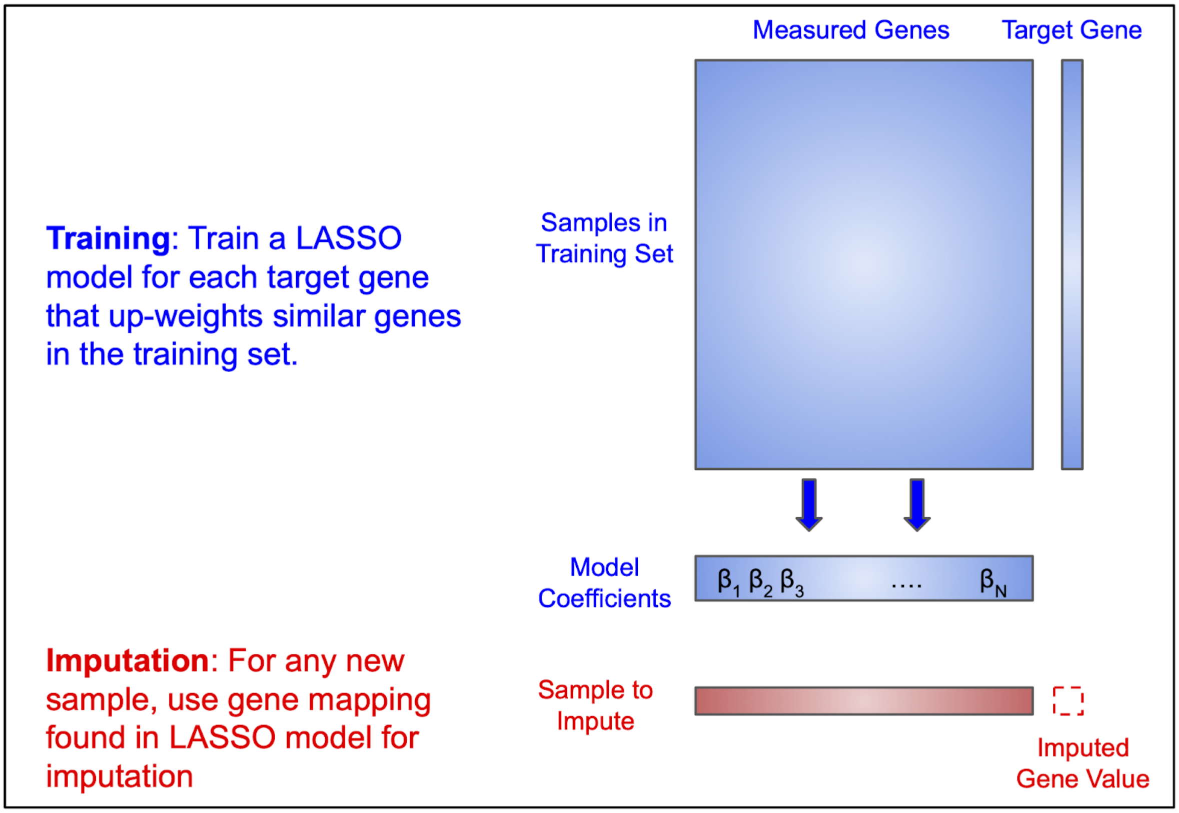


**Fig. S3. Schematic of *GeneLASSO*.** Blue color denotes training/fitting step and red color denotes imputation step.

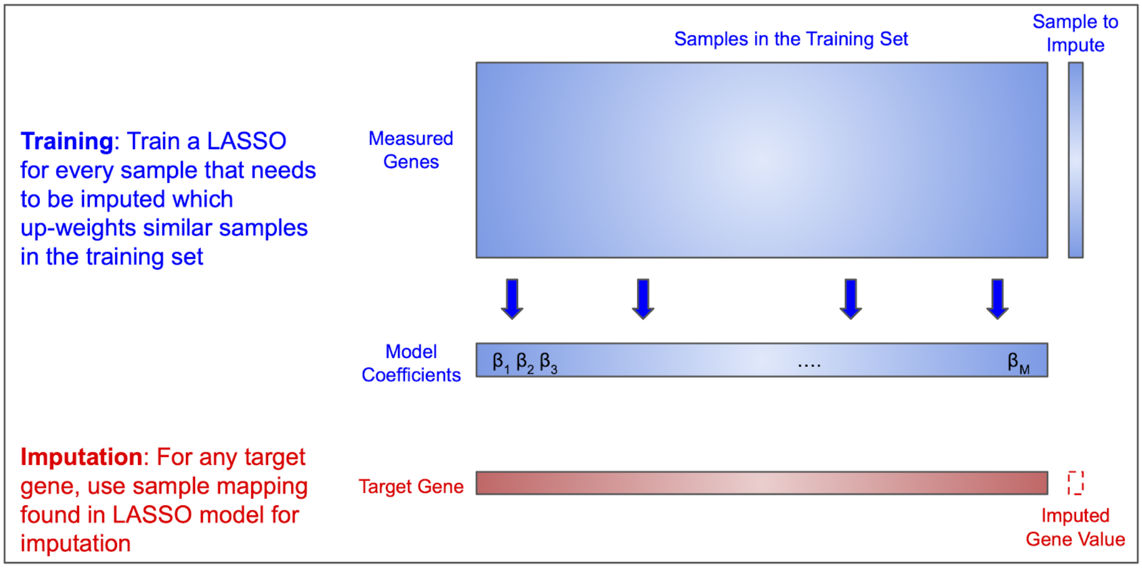


**Fig. S4. Schematic of *SampleLASSO*.** Blue color denotes training/fitting step and red color denotes imputation step.

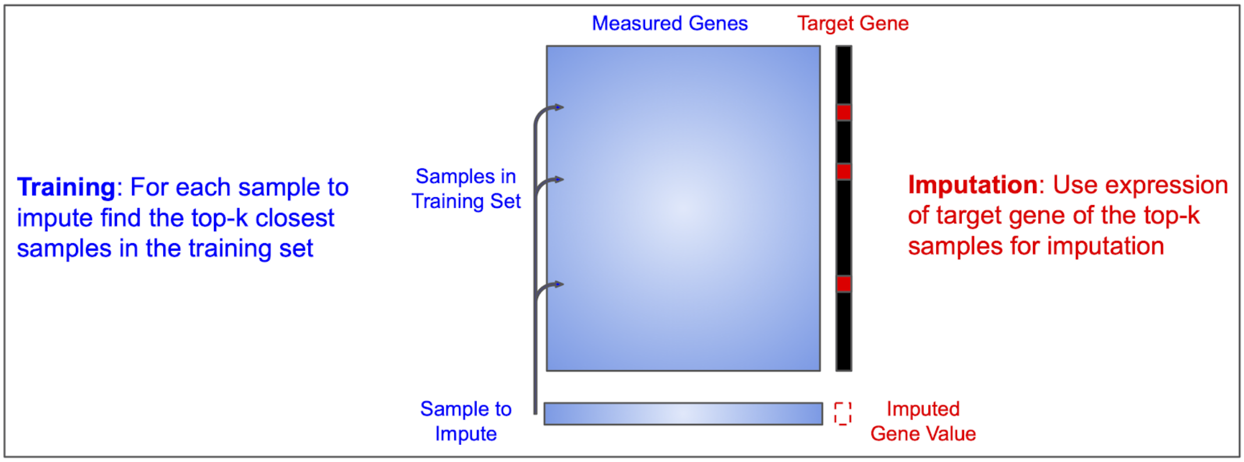


**Fig. S5. Schematic of *SampleKNN*.** Blue color denotes training/fitting step and red color denotes imputation step.

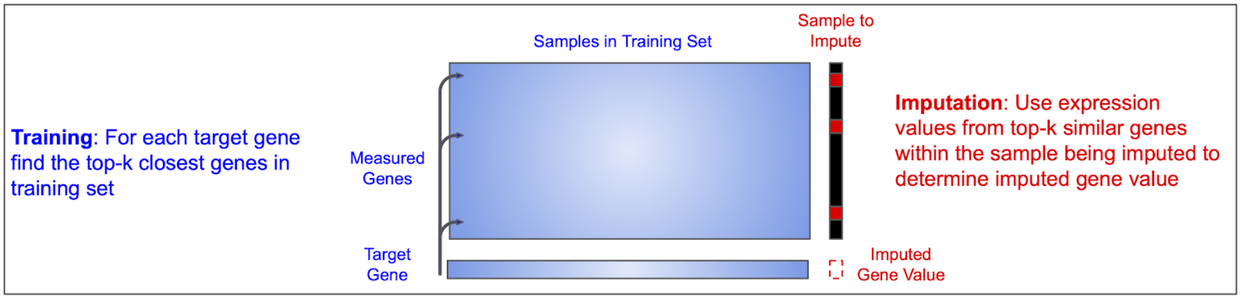


**Fig. S6. Schematic of *GeneKNN*.** Blue color denotes training/fitting step and red color denotes imputation step.

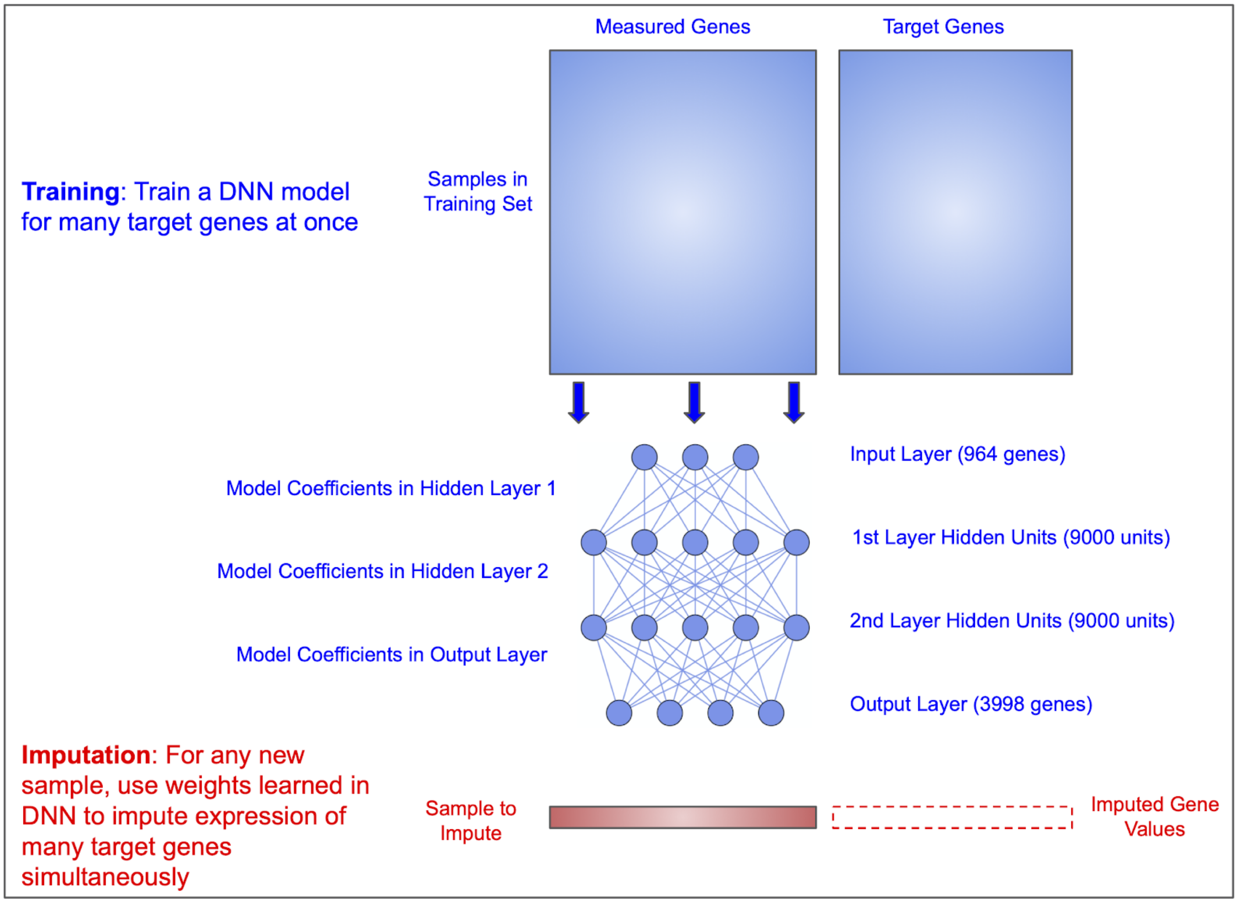


**Fig. S7. Schematic of *GeneDNN*.** Blue color denotes training/fitting step and red color denotes imputation step.

*Parameters for the best DNN*

The model parameters for the DNN used in this work were chosen based on those suggested in (Chen *et al.*, 2016). This includes using a dropout rate of 10%, Xavier Uniform weight initialization (Glorot and Bengio, 2010), a mini-batch size of 200 and 200 training epochs. We chose 2 hidden layers with 9000 units in each layer as this architecture was the best overall for doing same-technology and cross-technology imputation. It was not obvious what exact optimizer was used by (Chen *et al.*, 2016), so we performed hyperparameter tuning of the optimizer, using Adam (Kingma and Ba, 2017; Reddi *et al.*, 2018) and Adadelta (Zeiler, 2012). We implemented our DNN in *Keras* (Chollet, 2015) using a *Tensorflow* backend (Abadi *et al.*, 2016).

#### Section 1.4: Hyperparameter Tuning

Hyperparameter tuning was performed for all combinations of methods (SampleLASSO, GeneLASSO, SampleKNN, GeneKNN, GeneDNN), gene subsets (GPL96-570, LINCS) and tasks (Microarray-Microarray, RNAseq-Microarray, RNAseq-RNAseq). In LASSO methods, the hyperparameter that was tuned was the strength of the L1-regularization, referred to as alpha. In KNN methods, the hyperparameter that was tuned was the number of closest samples, referred to a k. In DNN methods, the hyperparameter that was tuned was the optimizer. The results of the hyperparameter tuning can be seen in Figs. S8-S12. Table 1 shows which hyperparameter yielded the optimal performance.

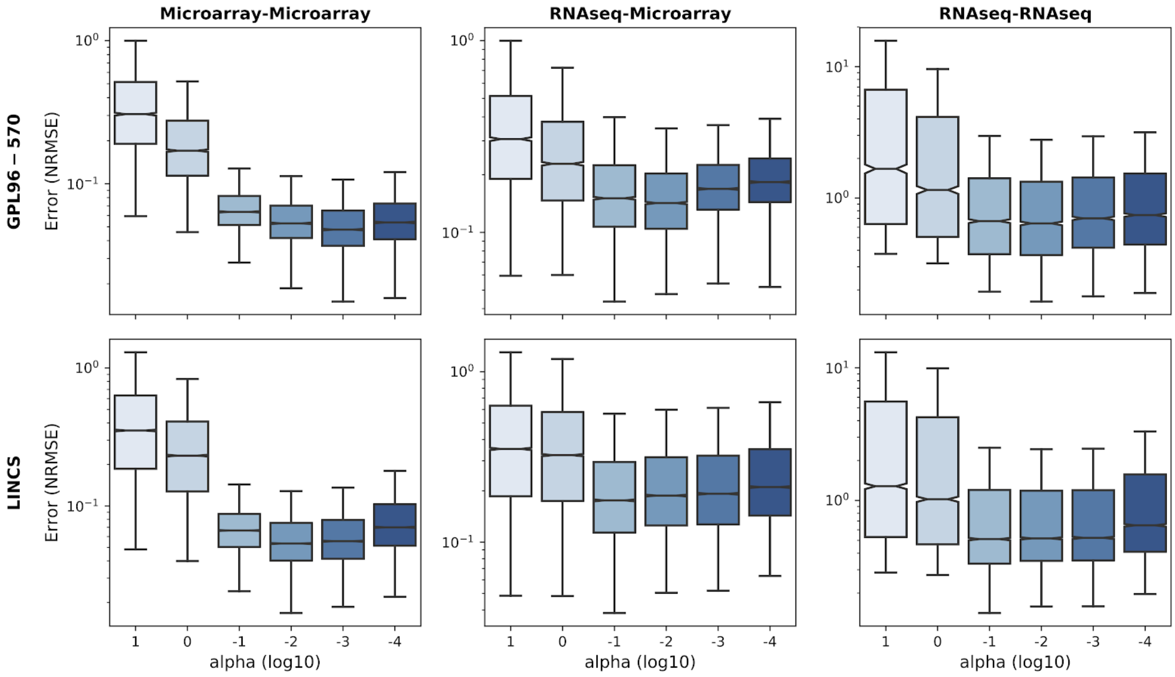


**Fig. S8. Hyperparameter tuning for *SampleLASSO*.** The rows are for the different gene subsets (top: GPL96-570 and bottom: LINCS). The columns are for the different technology combinations (left: Microarray-Microarray, middle: RNAseq-Microarray and right: RNAseq-RNAseq). The hyperparameter is alpha which controls the strength of the L1-regularization. The y-axis scale is log and the metric is NRMSE.

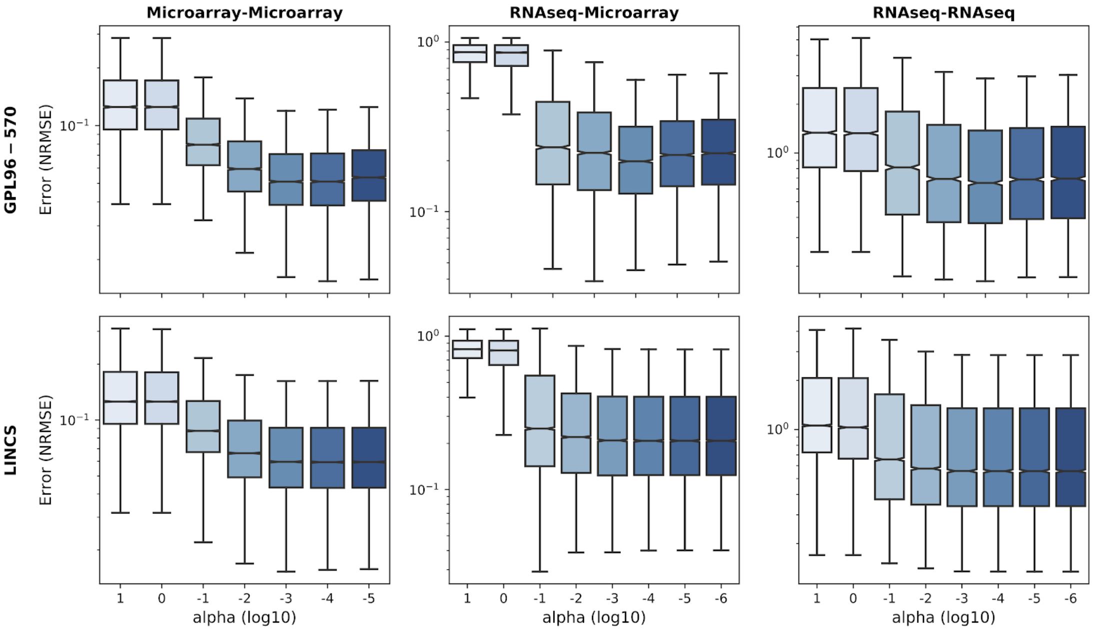


**Fig. S9. Hyperparameter tuning for *GeneLASSO*.** The rows are for the different gene subsets (top: GPL96-570 and bottom: LINCS). The columns are for the different technology combinations (left: Microarray-Microarray, middle: RNAseq-Microarray and right: RNAseq-RNAseq). The hyperparameter is alpha which controls the strength of the L1-regularization. The y-axis scale is log and the metric is NRMSE.

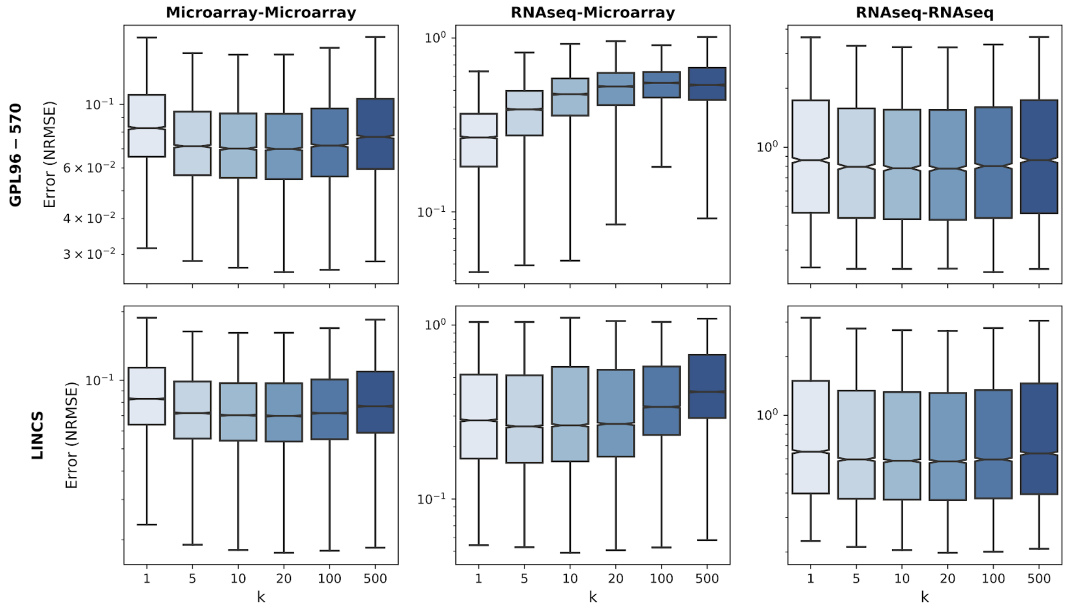


**Fig. S10. Hyperparameter tuning for *SampleKNN*.** The rows are for the different gene subsets (top: GPL96-570 and bottom: LINCS). The columns are for the different technology combinations (left: Microarray-Microarray, middle: RNAseq-Microarray and right: RNAseq-RNAseq). The hyperparameter is k which controls the number of closest examples to use.The y-axis scale is log and the metric is NRMSE.

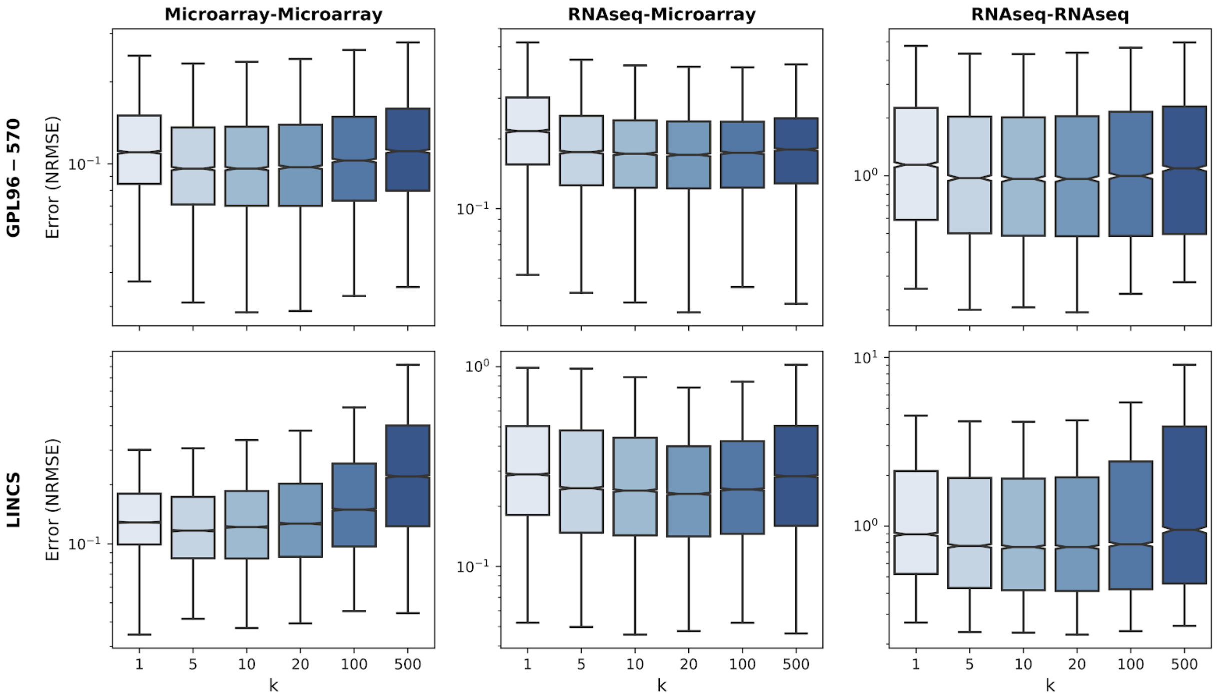


**Fig. S11. Hyperparameter tuning for *GeneKNN*.** The rows are for the different gene subsets (top: GPL96-570 and bottom: LINCS). The columns are for the different technology combinations (left: Microarray-Microarray, middle: RNAseq-Microarray and right: RNAseq-RNAseq). The hyperparameter is k which controls the number of closest examples to use. The y-axis scale is log and the metric is NRMSE.

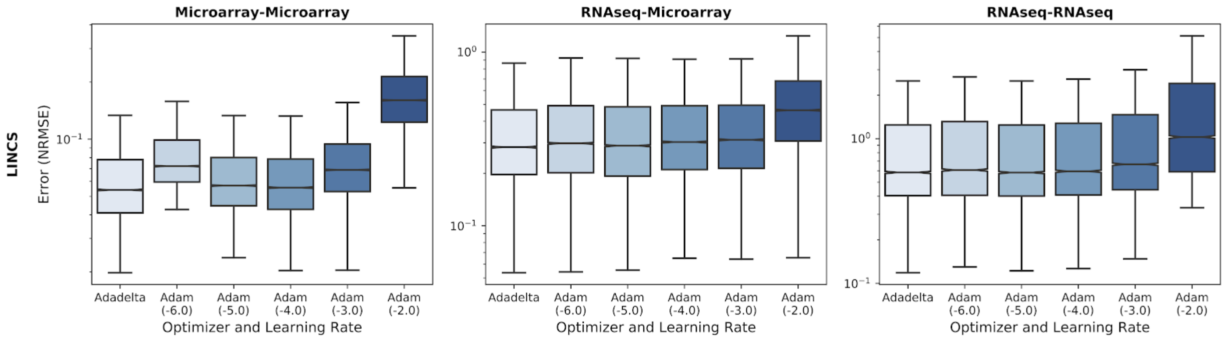


**Fig. S12. Hyperparameter tuning for *GeneDNN*.** The columns are for the different technology combinations (left: Microarray-Microarray, middle: RNAseq-Microarray and right: RNAseq-RNAseq). GeneDNN was only implemented for the LINCS gene subset. The hyperparameter is the optimizer and the log10 of the learning rate (Adadelta doesn't require an initial learning rate). These results show the error using the full validation set after the best model was selected across all epochs of training using the ModelCalback function in Keras. The y-axis scale is log and the metric is NRMSE.

**Table S1: Best Models Settings from Hyperparameter Tuning.** The NRMSE value is the median NRMSE value across all genes for that hyperparameter set.

| | | Method | Model Setting | Optimal Choice | NRMSE |
|---|---|---|---|---|---|
| **Microarray-Microarray** | **GPL96-570** | GeneKNN | k | 5.0 | 0.0961 |
| | **GPL96-570** | GeneLASSO | alpha | 0.001 | 0.0510 |
| | **GPL96-570** | SampleKNN | k | 20.0 | 0.0699 |
| | **GPL96-570** | SampleLASSO | alpha | 0.001 | 0.0478 |
| | **LINCS** | GeneDNN | optimizer | Adadelta | 0.0540 |
| | **LINCS** | GeneKNN | k | 5.0 | 0.1167 |
| | **LINCS** | GeneLASSO | alpha | 0.0001 | 0.0593 |
| | **LINCS** | SampleKNN | k | 20.0 | 0.0698 |
| | **LINCS** | SampleLASSO | alpha | 0.01 | 0.0534 |
| **RNAseq-Microarray** | **GPL96-570** | GeneKNN | k | 20.0 | 0.1708 |
| | **GPL96-570** | GeneLASSO | alpha | 0.001 | 0.1980 |
| | **GPL96-570** | SampleKNN | k | 1.0 | 0.2677 |
| | **GPL96-570** | SampleLASSO | alpha | 0.01 | 0.1422 |
| | **LINCS** | GeneDNN | optimizer | Adadelta | 0.2837 |
| | **LINCS** | GeneKNN | k | 20.0 | 0.2309 |
| | **LINCS** | GeneLASSO | alpha | 0.0001 | 0.2078 |
| | **LINCS** | SampleKNN | k | 5.0 | 0.2612 |
| | **LINCS** | SampleLASSO | alpha | 0.1 | 0.1757 |
| **RNAseq-RNAseq** | **GPL96-570** | GeneKNN | k | 20.0 | 0.9632 |
| | **GPL96-570** | GeneLASSO | alpha | 0.001 | 0.6533 |
| | **GPL96-570** | SampleKNN | k | 20.0 | 0.7792 |
| | **GPL96-570** | SampleLASSO | alpha | 0.01 | 0.6410 |
| | **LINCS** | GeneDNN | optimizer | Adam (1e-05) | 0.5833 |
| | **LINCS** | GeneKNN | k | 20.0 | 0.7506 |
| | **LINCS** | GeneLASSO | alpha | 1e-05 | 0.5560 |
| | **LINCS** | SampleKNN | k | 20.0 | 0.5811 |
| | **LINCS** | SampleLASSO | alpha | 0.1 | 0.5107 |

### Section 2: Supplemental Results

#### Section 2.1: Detailed Information on Method Comparisons

**Table S2: Detailed information on comparing the methods.** Percent SL Better is the percentage of times *SampleLASSO* is better than the other method. Log2 Effect Size is the log2 increase of *SampleLASSO* over the other method considering just the median value (a positive value is when *SampleLASSO* is the better performing method). P-Value is the significance between *SampleLASSO* and the other method based on a Wilcoxon rank-sum test.

| | | Median | Percent SL Better | Log2 Effect Size | P-Value |
|---|---|---|---|---|---|
| **Microarray-Microarray-GPL96-570** | **SampleLASSO** | 0.047 | N/A | N/A | N/A |
| | **GeneLASSO** | 0.050 | 0.91 | 0.088 | 0.00e+00 |
| | **SampleKNN** | 0.069 | 1.00 | 0.542 | 0.00e+00 |
| | **GeneKNN** | 0.096 | 1.00 | 1.024 | 0.00e+00 |
| **Microarray-Microarray-LINCS** | **SampleLASSO** | 0.053 | N/A | N/A | N/A |
| | **GeneDNN** | 0.054 | 0.76 | 0.031 | 0.00e+00 |
| | **GeneLASSO** | 0.058 | 0.91 | 0.137 | 0.00e+00 |
| | **SampleKNN** | 0.069 | 0.98 | 0.373 | 0.00e+00 |
| | **GeneKNN** | 0.113 | 1.00 | 1.094 | 0.00e+00 |
| **RNAseq-Microarray-GPL96-570** | **SampleLASSO** | 0.144 | N/A | N/A | N/A |
| | **GeneKNN** | 0.172 | 0.70 | 0.254 | 4.38e-223 |
| | **GeneLASSO** | 0.199 | 0.71 | 0.467 | 2.21e-299 |
| | **SampleKNN** | 0.265 | 0.80 | 0.881 | 0.00e+00 |
| **RNAseq-Microarray-LINCS** | **SampleLASSO** | 0.175 | N/A | N/A | N/A |
| | **GeneLASSO** | 0.208 | 0.62 | 0.251 | 0.00e+00 |
| | **GeneKNN** | 0.228 | 0.84 | 0.384 | 0.00e+00 |
| | **SampleKNN** | 0.263 | 0.76 | 0.589 | 0.00e+00 |
| | **GeneDNN** | 0.291 | 0.77 | 0.734 | 0.00e+00 |
| **RNAseq-RNAseq-GPL96-570** | **SampleLASSO** | 0.866 | N/A | N/A | N/A |
| | **GeneLASSO** | 0.871 | 0.41 | 0.009 | 6.96e-03 |
| | **SampleKNN** | 0.993 | 0.93 | 0.198 | 0.00e+00 |
| | **GeneKNN** | 1.133 | 0.98 | 0.388 | 0.00e+00 |
| **RNAseq-RNAseq-LINCS** | **GeneDNN** | 0.693 | 0.18 | -0.021 | 0.00e+00 |
| | **SampleLASSO** | 0.703 | N/A | N/A | N/A |
| | **GeneLASSO** | 0.744 | 0.62 | 0.082 | 0.00e+00 |
| | **SampleKNN** | 0.777 | 0.84 | 0.145 | 0.00e+00 |
| | **GeneKNN** | 0.914 | 1.00 | 0.379 | 0.00e+00 |

#### Section 2.2: Results for Spearman and MAE metrics

In this section, we present the results in terms of the spearman correlation and the mean absolute error (MAE). The spearman correlation for a gene, $(g_i)$, is given by

$$Spearman(g_i) = \frac{cov(\hat{x}_R, x_R)}{\sigma_{\hat{x}_R}\sigma_{x_R}} \qquad \text{eqn. S1}$$

where $\hat{x}_R$ and $x_R$ are the imputed and real values for all the samples converted to ranks, respectively, $cov$ is the covariance and $\sigma$ is the standard deviation. The mean absolute error for a gene, $(g_i)$, is given by

$$MAE(g_i) = \sum_{j=1}^{S} \left| \hat{g}_{i,j} - g_{i,j} \right| / S \qquad \text{eqn. S2}$$

Where $S$ is the number of samples, and $\hat{g}_{i,j}$, $g_{i,j}$ are the imputed and real expression values, respectively, for the $i^{th}$ gene in the $j^{th}$ sample.

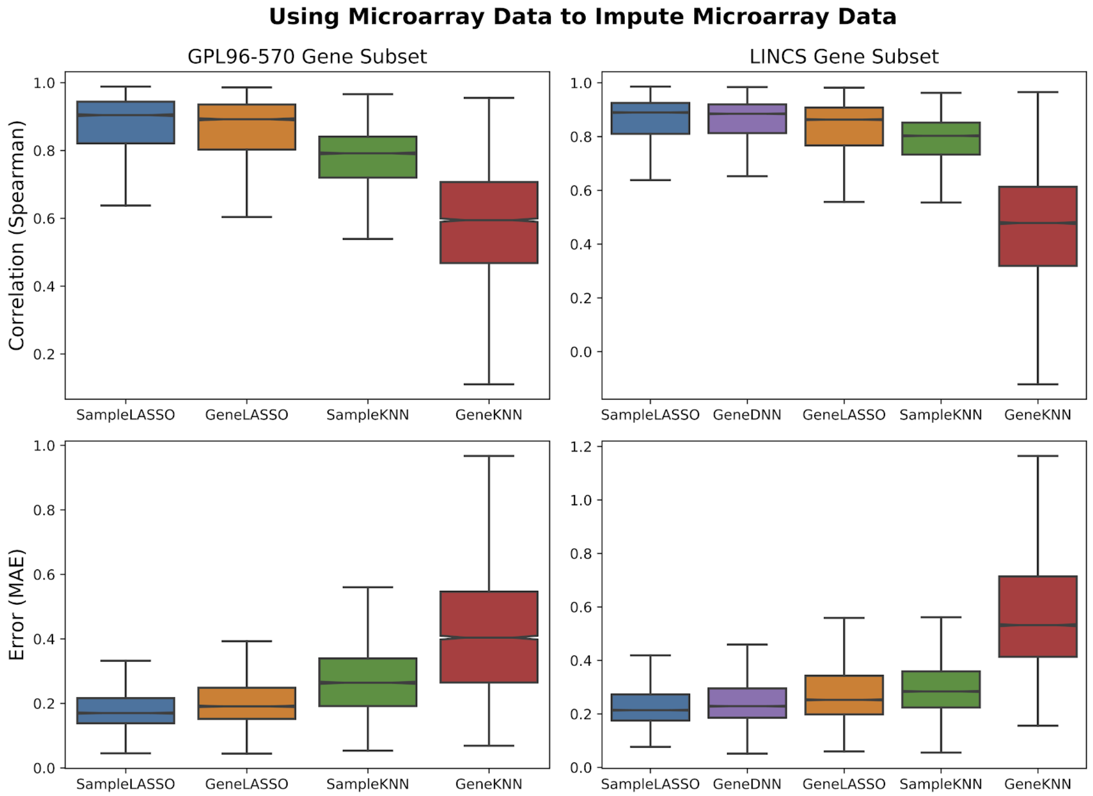


**Fig S13. Performance of imputation models on microarray data with Spearman and MAE metrics.** The performance of the five methods imputation models (*SampleLASSO*, *GeneDNN*, *GeneLASSO*, *SampleKNN*, and *GeneKNN*) are compared for using microarray data to impute microarray data.

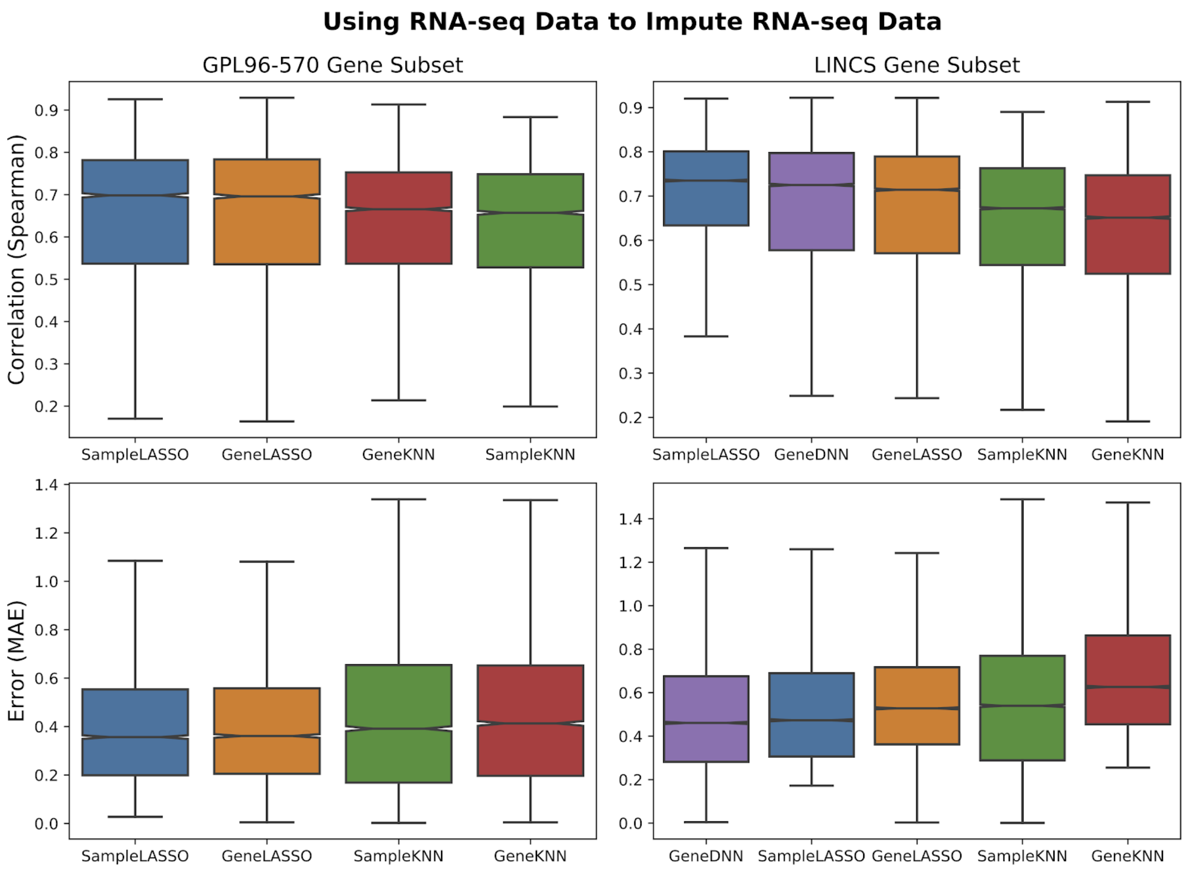


**Fig S14. Performance of imputation models on RNA-seq data with Spearman and MAE metrics.** The performance of the five methods imputation models (*SampleLASSO*, *GeneDNN*, *GeneLASSO*, *SampleKNN*, and *GeneKNN*) are compared for using RNA-seq data to impute RNA-seq data.

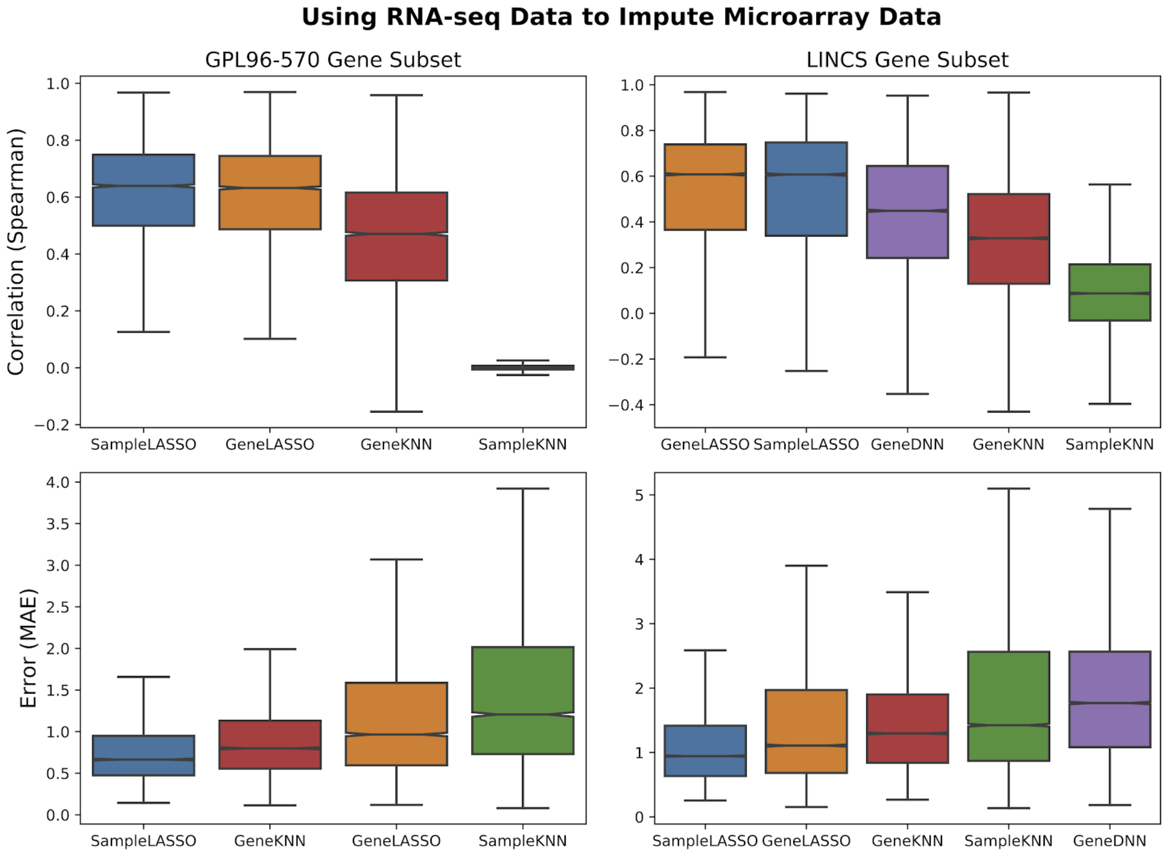


**Fig S15. Performance of imputation methods for cross-technology imputation with Spearman and MAE metrics.** The performance of the five methods imputation models (*SampleLASSO*, *GeneDNN*, *GeneLASSO*, *SampleKNN*, and *GeneKNN*) are compared for using RNA-seq data to impute microarray data.

#### Section 2.3: Results for RNA-seq to RNA-seq

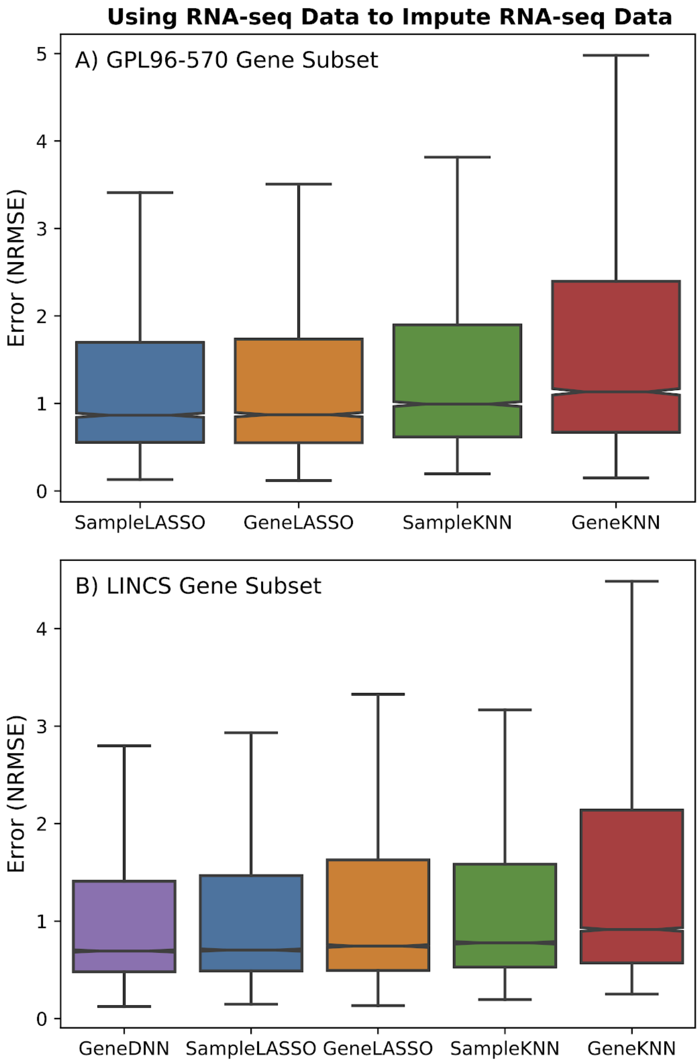


**Fig S16. Performance of imputation methods on RNA-seq Data.** The performance of the five imputation methods (*SampleLASSO*, *GeneDNN*, *GeneLASSO*, *SampleKNN*, *GeneKNN*) are compared across two gene subsets (GPL96-570 and LINCS) using RNA-seq data to impute RNA-seq data. The evaluation metric is NRMSE.

#### Section 2.4: Evaluations considering the expression levels and variance

In this section, we analyze how the performance of the methods changes for two gene properties; the mean expression of a gene and the variance of the expression. Genes were split up into low, medium and high bins for each property, and each panel in a figure [Figs. S17-S22] is the intersection of genes included in the two bins. See the plotting notebook in the associated

GitHub repo (https://github.com/krishnanlab/Expresto) for the breakdown of mean and variance values in each bin as well as the number of points in each boxplot.

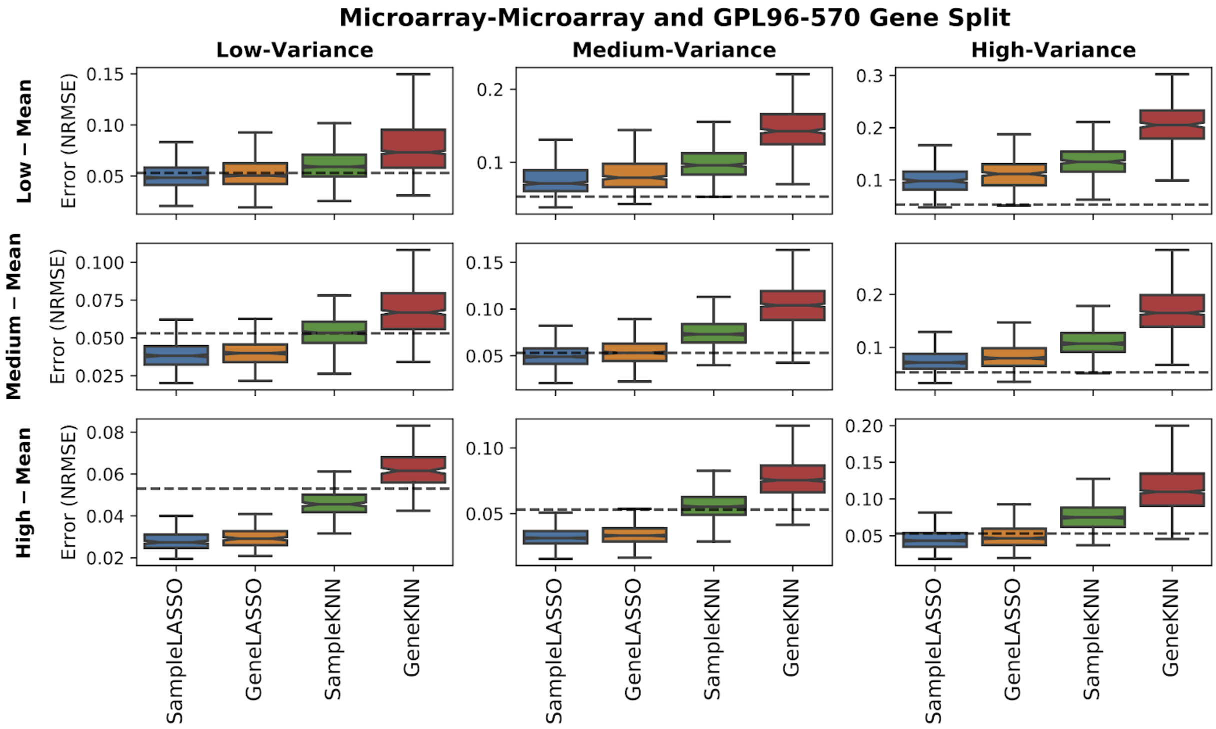


**Fig. S17. Results broken up by mean and variance of gene expression for using microarray data to impute microarray data for the GPL96-570 gene subset.** The dotted line is the median value when considering all genes for *SampleLASSO* (this is to help compare performances across the panels).

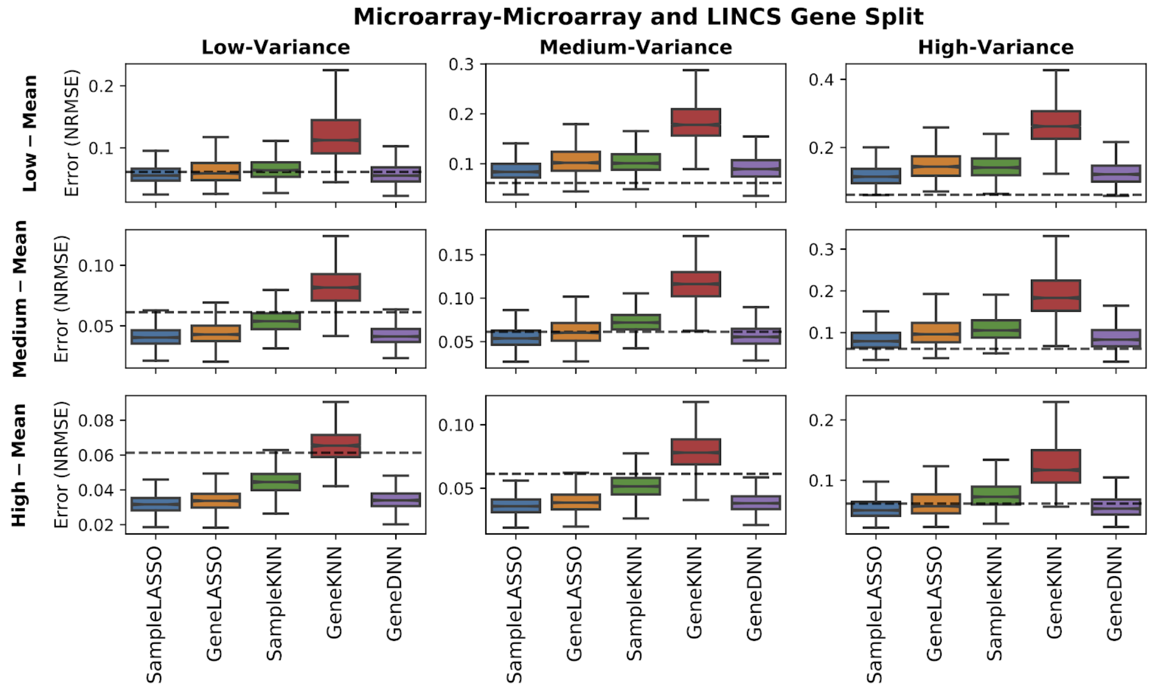


**Fig. S18. Results broken up by mean and variance of gene expression for using microarray data to impute microarray data for the LINCS gene subset.** The dotted line is the median value when considering all genes for *SampleLASSO* (this is to help compare performances across the panels).

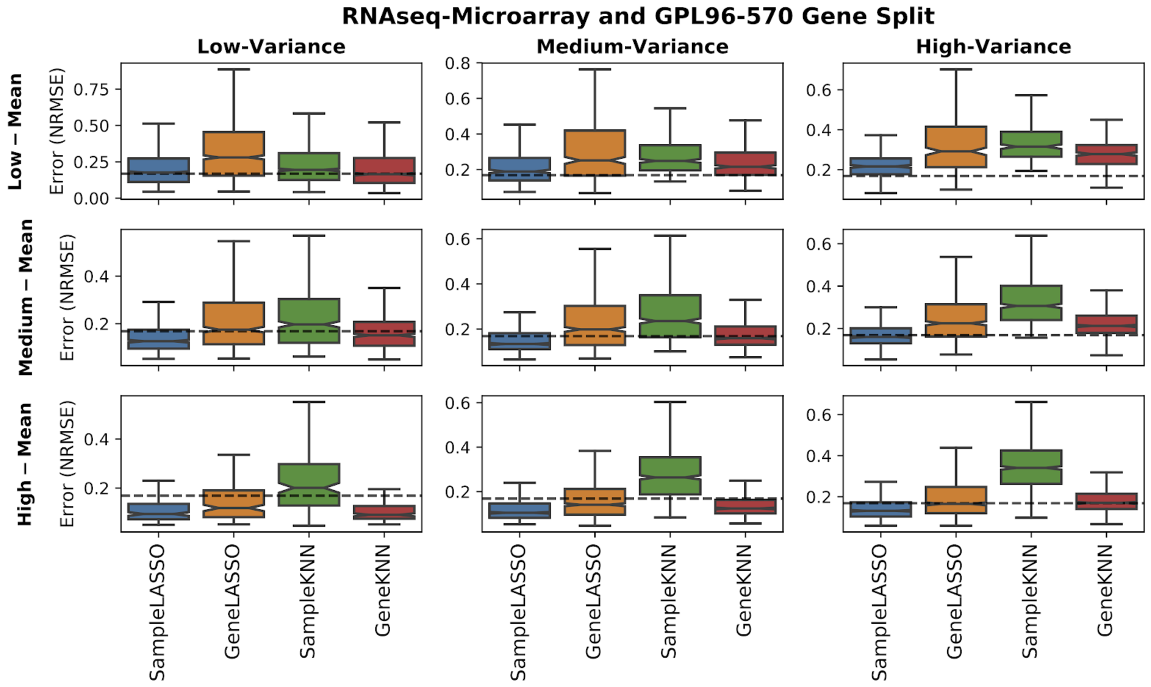


**Fig. S19. Results broken up by mean and variance of gene expression for using RNA-seq data to impute microarray data for the GPL96-570 gene subset.** The dotted line is the median value when considering all genes for *SampleLASSO* (this is to help compare performances across the panels).

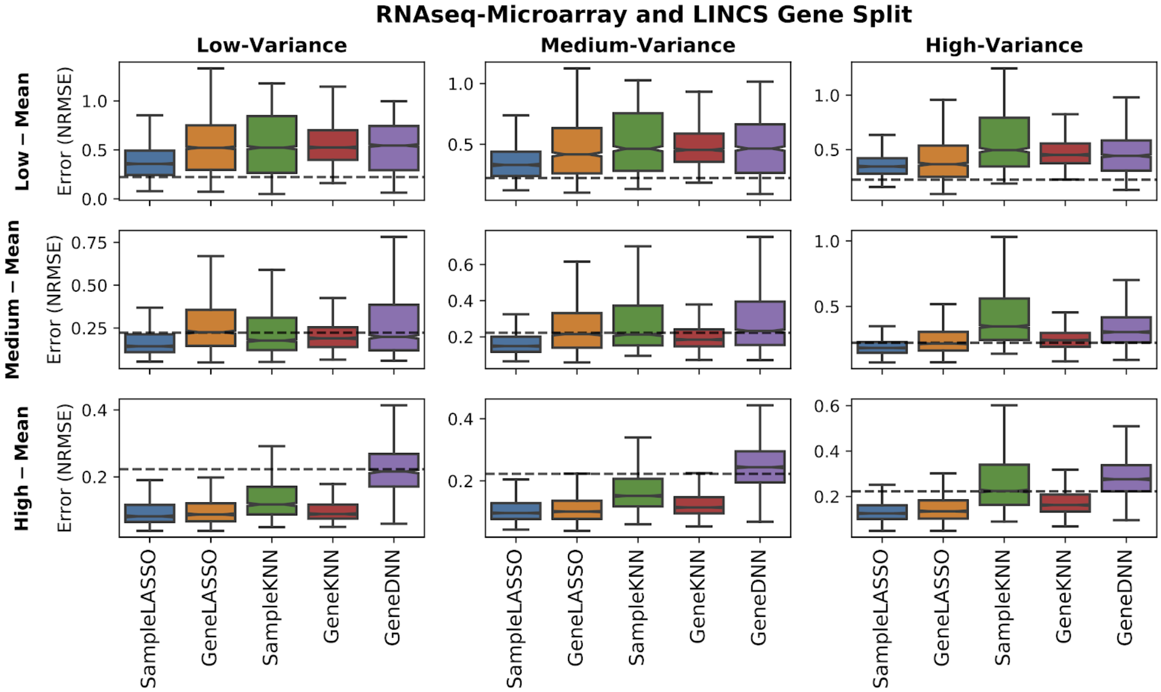


**Fig. S20. Results broken up by mean and variance of gene expression for using RNA-seq data to impute microarray data for the LINCS gene subset.** The dotted line is the median value when considering all genes for *SampleLASSO* (this is to help compare performances across the panels).

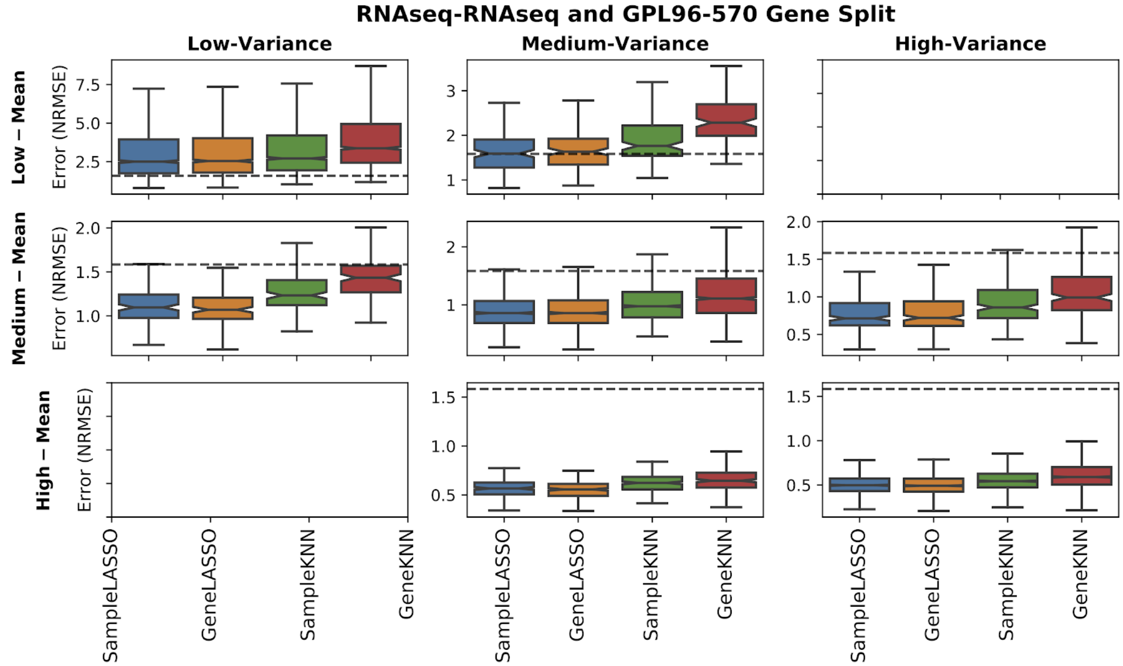


**Fig. S21. Results broken up by mean and variance of gene expression for using RNA-seq data to impute RNA-seq data for the GPL96-570 gene subset.** The dotted line is the median value when considering all genes for *SampleLASSO* (this is to help compare performances across the panels).

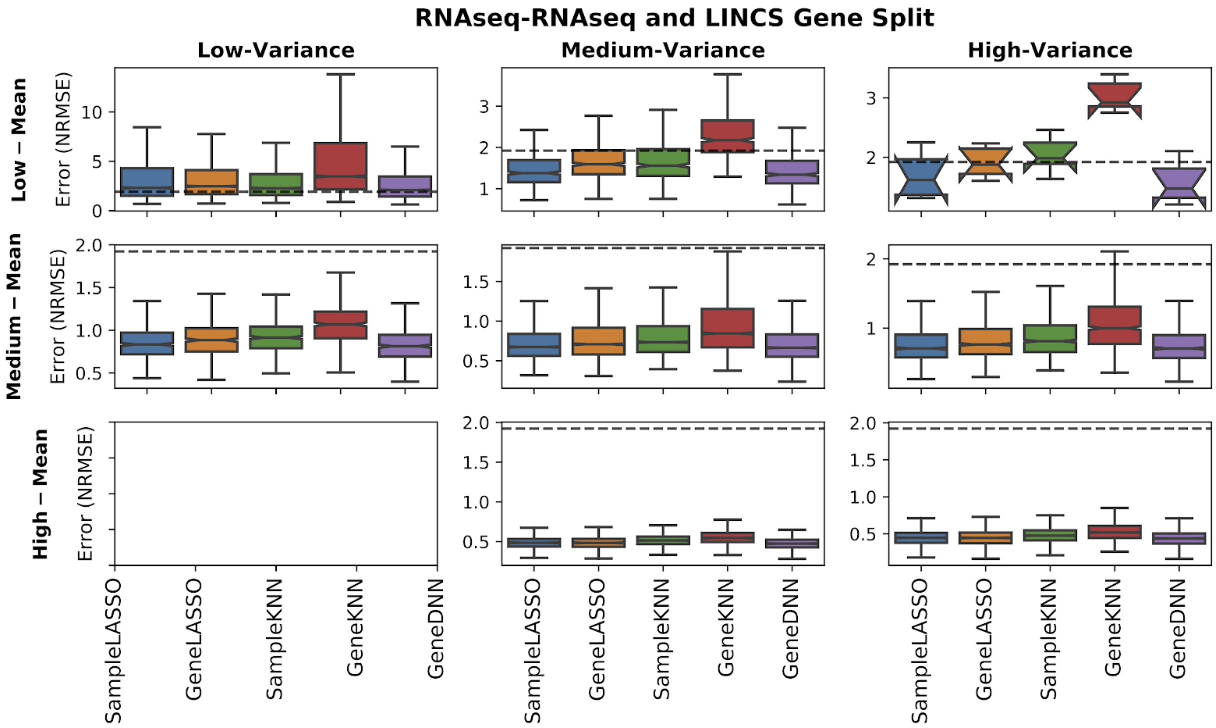


**Fig. S22. Results broken up by mean and variance of gene expression for using RNA-seq data to impute RNA-seq data for the LINCS gene subset.** The dotted line is the median value when considering all genes for *SampleLASSO* (this is to help compare performances across the panels).

#### Section 2.5: Supplemental Material for Beta Analysis

The breakdown of the number of how many samples for each tissue were used in the beta-coefficient analysis can be found in Table S3. We additionally analyzed the beta-coefficients by grouping the z-scores of all non-target tissue samples together into a non-target group, as well as grouping the z-scores for all target tissue samples together. This was done for each tissue separately [Fig. S23]. An example of information returned by the Expresto software (https://github.com/krishnanlab/Expresto) can be seen in Table S4.

**Table S3. Statistics of Data Used in Beta-Coefficient Analysis**

| | Test Set | | Training Set | |
|---|---|---|---|---|
| **Tissue** | **Number of GSMs** | **Number of GSEs** | **Number of GSMs** | **Number of GSEs** |
| Blood | 63 | 5 | 1197 | 33 |
| Liver | 40 | 6 | 776 | 27 |
| Breast | 40 | 6 | 770 | 25 |
| Brain | 39 | 3 | 752 | 19 |
| Lung | 29 | 6 | 662 | 10 |
| Kidney | 11 | 3 | 240 | 6 |

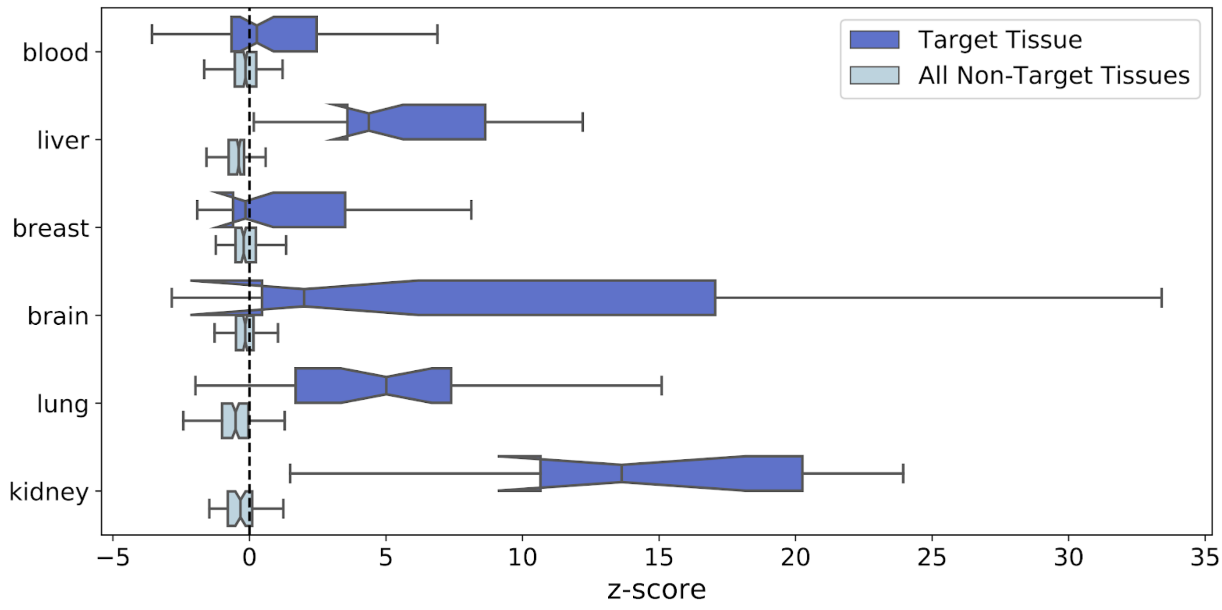


**Fig. S23. Model interpretability of the target tissue versus rest of tissues combined.** For each test sample labeled for a given tissue, we found the z-score for all six tissues considered. The boxplots show the distribution of z-scores for the target tissue, as well as all non-target tissues.

**Table S4. Example of information that can be obtained from the user_function in the *Expresto* software released with this work**

| **Target sample being imputed** | | **Training samples with the three highest model coefficients** | | | |
|---|---|---|---|---|---|
| | | | **First** | **Second** | **Third** |
| **Sample**<br>**Study**<br>**Annotation** | GSM478457<br>GSE19279/GSE19281<br>Title: Normal pancreas, 3 (U133A) | **Sample**<br>**Study**<br>**Beta-coeff**<br>**Annotation** | GSM175950<br>GSE7307<br>0.34<br>Title: Pancreas SG1 Normal | GSM388108<br>GSE15471<br>0.15<br>Source name: pancreas | GSM388111<br>GSE15471<br>0.14<br>Source name: pancreas |
| **Sample**<br>**Study**<br>**Annotation** | GSM664048<br>GSE26971<br>Source name: primary breast cancer sample, fresh-frozen | **Sample**<br>**Study**<br>**Beta-coeff**<br>**Annotation** | GSM102499<br>GSE2109<br>0.15<br>Title: Breast - 129692 | GSM151315<br>GSE6532<br>0.11<br>Source name: breast | GSM687049<br>GSE27830/GSE54219<br>0.09<br>Title: primary breast cancer, sample_1927 |
| **Sample**<br>**Study**<br>**Annotation** | GSM4005<br>GSE475<br>Source name: Human diaphragm | **Sample**<br>**Study**<br>**Beta-coeff**<br>**Annotation** | GSM342677<br>GSE13070<br>0.15<br>Source name: skeletal muscle (vastus lateralis) | GSM42736<br>GSE2328<br>0.13<br>Source name: skeletal muscle | GSM342884<br>GSE13070<br>0.12<br>Source name: skeletal muscle (vastus lateralis) |

#### Section 2.6: Loss Curves for GeneDNN

In this section, we present the loss curves for the DNNs [Figs. S24-S26]. For the DNNs the unmeasured genes were broken into four sets, and in the figures each panel represents a model trained on each of these four sets. The data was generated using the *csv_logger* callback in *Keras* using the mean absolute error across all examples of the output of the mode. For all plots the training error displays the expected trends. Two interesting observations are for using RNA-seq data to impute RNA-seq data [Fig. S25], the validation data has a lower loss than the training data. When using RNA-seq data to impute microarray data [Fig. S26], the validation loss is much higher than the training loss, suggesting that the model is overfitting to the training data.

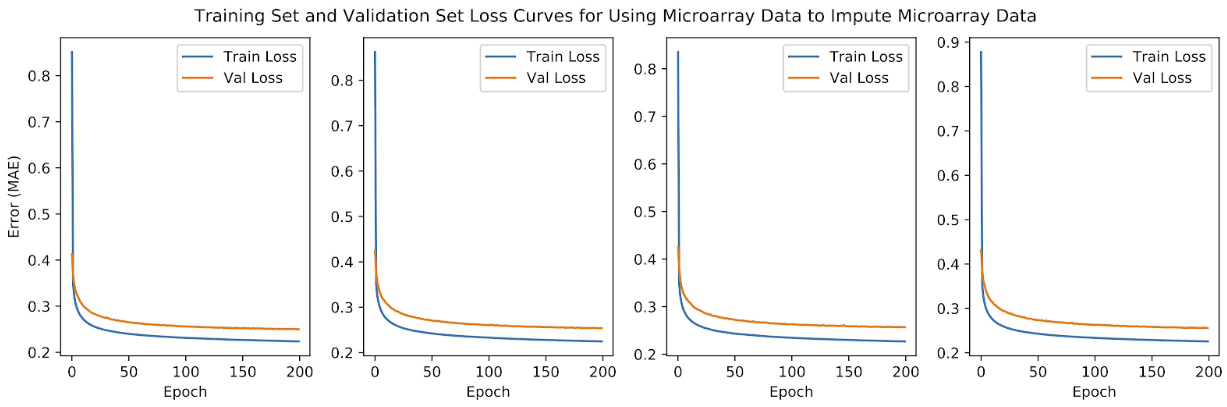


**Fig S24. DNN loss curves for using microarray data to impute microarray data.**

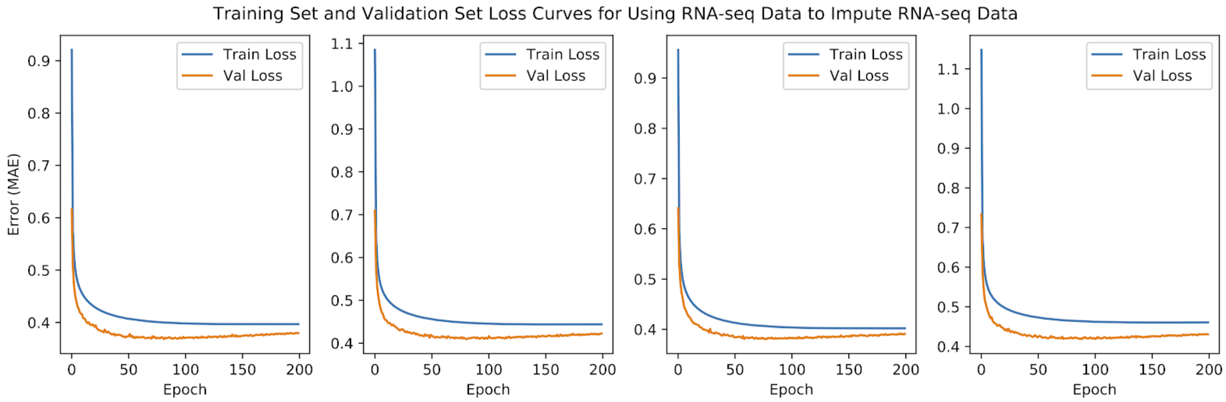


**Fig S25. DNN loss curves for using RNA-seq data to impute RNA-seq data.**

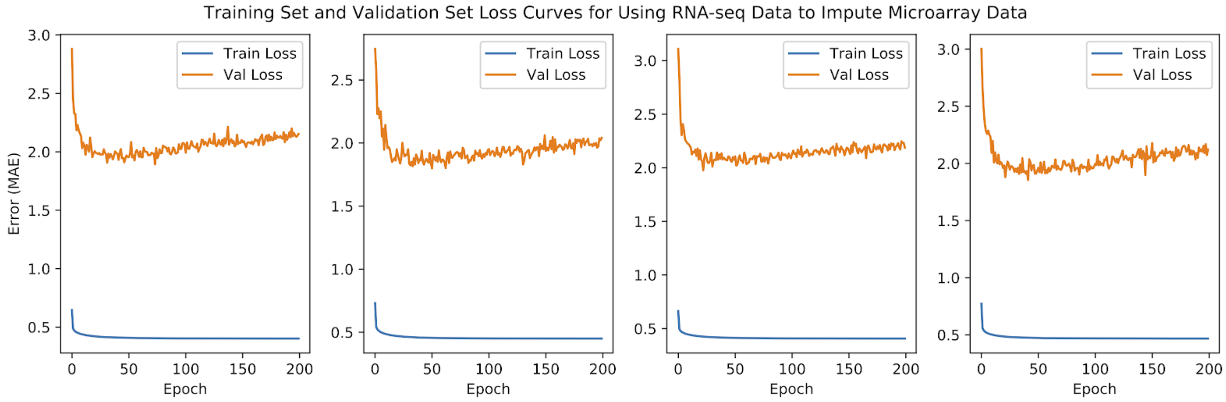


**Fig S26. DNN loss curves for using RNA-seq data to impute microarray data.**
